# Remodeling the central metabolism of *Escherichia coli* enables a universal chassis

**DOI:** 10.1101/2023.05.08.539912

**Authors:** Min Liu, Likun Guo, Meitong Huo, Xinjun Feng, Zhe Zhao, Qingsheng Qi, Mo Xian, Guang Zhao

## Abstract

*E. coli* is the host of choice to produce a wide variety of chemicals and proteins. Overflow metabolism is considered as the widespread and major obstacle in microbial synthesis, and overcoming this common bottleneck may enable a universal chassis. Here, we constructed an *E. coli* universal chassis (ABKS strain) with significantly suppressed overflow metabolism, presenting similar growth rate, decreased glucose consumption, and increased production of desired chemicals and proteins when compared with wild-type BL21(DE3) strain. Furthermore, we demonstrated that metabolic flux of ABKS strain was reprogrammed from TCA cycle to glyoxylate bypass at isocitrate node via the synergistic effect of multi-layer regulation in gene transcription and protein modification. This metabolic reconfiguration alleviates overflow metabolism, avoids CO_2_ release in TCA cycle, finally improving the carbon atom economy in bioprocess. Our chassis has widespread and practical use for elevating the production and yield of multiple desired chemicals and proteins from different carbon source. The metabolic reconfiguration also provides theoretical basis for rational design of efficient bioproduction strains.

## Introduction

In recent years, the bioproduction of value-added chemicals and materials is getting momentum, and is believed to be a substitute of unsustainable fossil-based chemical processes in the future. *Escherichia coli* is one of the host organisms of choice, and has been engineered successfully to produce a wide variety of compounds including fuels (*1, 2*), plastic monomers and polymers (*3, 4*), amino acids (*5, 6*), aromatics (*7*), pigments (*8*), and so on. Yet again, the production of a large number of compounds has yet to successfully reached commercial scale (*9*). Improving titers and yields of biotransformation processes to meet industrial production requires rational manipulation of the selected host. At present, the conventional metabolic regulation is still aimed at optimizing the biosynthetic pathway of target chemical, producing an engineering strain with limited application, although the process is very time-consuming and labor-intensive. The efficient bioproduction is not only relied on the biosynthetic pathway, but also influenced deeply by the endogenous metabolic network of the host cells. Therefore, it is hoped to generate a universal chassis with efficient bioproduction through overcoming a widespread obstacle in the endogenous metabolic network.

When growing on glucose as sole carbon source aerobically, *E. coli* uptakes and metabolizes glucose rapidly, and accumulates a large amount of acetate in the culture broth, which is well-documented and was defined as overflow metabolism (*10*). Acetate accumulation accounts for 10 to 30% of carbon flux from glucose under fully aerated conditions (*11*), resulting in reduction of biomass and product yield, and compromise of the economy of industrial bioproduction (*12, 13*). Overflow metabolism is considered as one of the major obstacles in *E. coli*’s bioprocess, and has attracted the attentions of biotechnologists and microbiologists for over four decades (*10, 12, 14, 15*). Thus, metabolic reconfiguration targeting overflow metabolism may enable a universal chassis with improved bioproduction of multiple desired chemicals and proteins.

The causes of overflow metabolism remain incompletely understood. It is generally accepted that the excretion of those fermentation products is caused by the imbalance between rapid glucose catabolism and the limited respiratory capacity of *E. coli* (*10, 12*). A large amount of reducing equivalents produced in glycolysis pathway and tricarboxylic acid (TCA) cycle cannot be oxidized completely in the electron transfer process of respiratory chain. The increase of intracellular NAD(P)H concentration further inhibits the activities of pyruvate dehydrogenase (*16*) and citrate synthetase (*17*), and activates the TCA cycle negative regulator ArcA (*18, 19*), leading to intracellular accumulation of pyruvate and acetyl-CoA, and insufficient supply of ATP and NAD^+^. Given the circumstances, acetyl-CoA is converted into acetate through the PTA-AckA pathway in coupling with generation of one ATP molecule (*20*), and pyruvate is reduced into lactate by lactate dehydrogenase LdhA to recycle NAD^+^ (*21, 22*). In fact, lactate was also reported as a major byproduct besides acetate under these conditions (*23, 24*). So, acetate production and lactate production are joint processes responding to *E. coli* reaching the maximum oxygen consumption rate when growing on glucose aerobically.

Many attempts have been made to repress overflow metabolism, and some improvements have been achieved through deleting the byproducts related genes *ackA* and *ldhA* (*25, 26*), overexpressing the acetate assimilation gene *acs* (*27*), slowing down the glucose uptake by deletion of genes encoding glucose phosphotransferase system (PTS) (*28–30*), upregulating the expression level of TCA cycle genes by deletion of *arcA* gene (*31*), and providing outlet for the “excess” intracellular NAD(P)H by overexpression of exogenous NADH oxidase (*32*). However, it is hard for inactivation or overexpression of single gene to deal with the overflow metabolism problem and to improve the yield of target product to the level we desire because overflow metabolism is controlled by a regulatory network with a high level of complexity, integrating regulation mechanisms operating at different cellular levels (*33–35*).

In this study, we constructed a universal chassis ABKS with significantly suppressed overflow metabolism, similar growth rate, decreased glucose consumption and elevated production of desired chemicals and proteins when compared with BL21(DE3) wild-type strain. Analyses at multiple layers of gene transcription, metabolites, protein acetylation and activities of metabolic enzymes were carried out, and revealed that the central metabolic flux of the ABKS strain was remodeled from TCA cycle to glyoxylate bypass at isocitrate node, which alleviates the overflow metabolism, avoids the natural carbon loss in TCA cycle and improves the carbon economy. Given that, the ABKS strain, as a universal chassis, have great potential for elevating the production and yield of desired chemicals and proteins and will secure a place in industrial bioproduction.

## Results

### Construction of strains with suppressed overflow metabolism

To repress overflow metabolism, five genes related in central metabolism regulation and acetate production were selected as candidates for *E. coli* genetic modifications (Supplementary Fig. 1): (1) *arcA* gene, along with *arcB*, encodes a quinone-dependent two-component system involved in regulation of respiratory and fermentative metabolism (*36, 37*), and deletion of *arcA* will relieve the transcriptional repression of genes in TCA cycle and glyoxylate shunt (*32*). (2) *iclR* gene encodes a transcriptional repressor which binds pyruvate and represses the expression of *aceBAK* operon encoding glyoxylate shunt enzymes (*38*). (3) *csrB* gene encodes a non-coding small RNA which prevents the RNA-binding protein CsrA from binding target transcripts and alleviates CsrA-dependent translation inhibition. This Csr (carbon storage regulator) system of *E. coli* influences quite a few physiological processes including central carbon metabolism, biofilm development, motility, etc. (*39*) (4) acetyl-CoA synthase gene *acs* was overexpressed to enhance acetate utilization. (5) acetate kinase gene *ackA* was mutated to reduce the conversion of acetyl-CoA to acetate. All these five genes have proven helpful to improve the yields of biomass and target chemicals individually (*25, 27, 31, 40, 41*).

In *E. coli* BL21(DE3) strain, *arcA*, *iclR* and *ackA* were knocked out, the transcription of *csrB* was increased by insertion of a strong promoter P_T7_ ahead of this gene in chromosome, and *acs* gene was cloned into a plasmid under control of IPTG-induced T7 promoter. These genetic modifications were carried out individually or combined randomly, and total of 31 *E. coli* BL21(DE3) mutant strains were constructed. Among them, two mutants, AIS (Δ*arcA* Δ*iclR* p*acs*) and ABS (Δ*arcA* P_T7_-*csrB* p*acs*), were omitted from further characterization due to their severely impaired growth. Phloroglucinol (PG) and 3-hydroxypropionate (3HP) are both important platform chemicals which have been successfully synthesized in our previous studies (*4, 42*), and were adapted as a proxy to evaluate the strains constructed here. So, *E. coli* BL21(DE3) wild-type strain and other 29 strains were introduced with the biosynthetic pathways for PG and 3HP respectively, and the resultant strains were applied to shaking-flask cultivation to determine their growth, glucose consumption and the productions of PG, 3HP, acetate and lactate.

Modifications of these five genes both alone and in combination could repress the overflow metabolism and improve the yields of PG and 3HP from glucose in various degrees (Fig. 1). Among them, a quadruple mutant ABKS (Δ*arcA* Δ*ackA* P_T7_-*csrB* p*acs*) represented the highest productions and yields of PG and 3HP. In ABKS strain, PG and 3HP were accumulated to concentrations of 2.79 ± 0.05 g/L and 4.42 ± 0.15 g/L respectively, while BL21(DE3) wild-type strain only produced 0.44 ± 0.01 g/L PG and 2.29 ± 0.05 g/L 3HP, and the yields of PG and 3HP from glucose in ABKS strain were 5.43- and 2.25-time higher than those in wild-type strain.

**Fig. 1.**
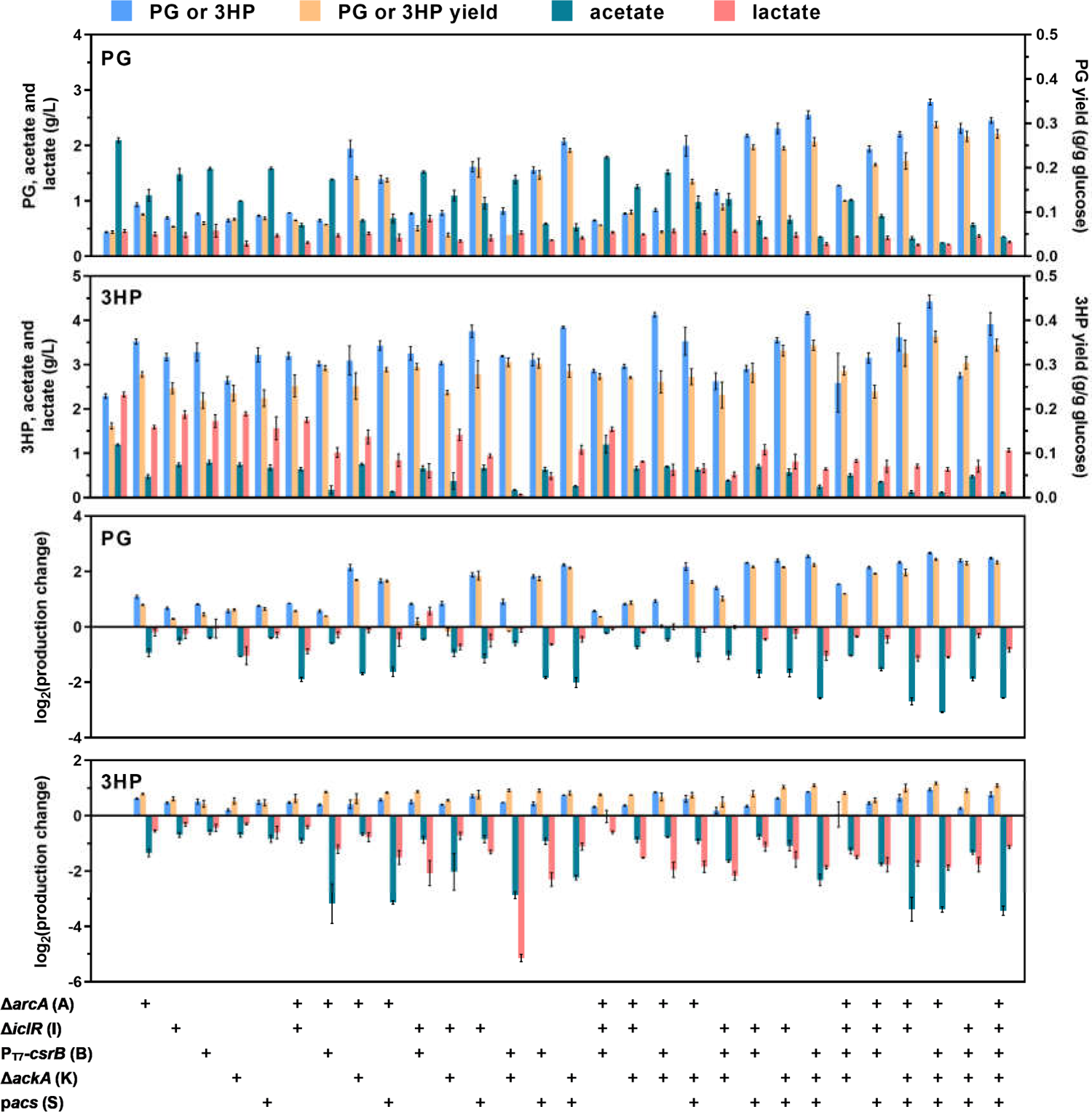
Effects of genetic modifications on productions of target chemicals and byproducts. Productions of phloroglucinol (PG), 3-hydroxypropionate (3HP), acetate and lactate by *E. coli* BL21(DE3) wild-type strain and 29 genetically modified strains were shown here and in Supplementary Table 1. The biosynthetic pathways for PG and 3HP were introduced into these strains respectively, and the resultant strains were applied to shaking-flask cultivation. Production change = production of genetically modified strain/production of wild-type strain. *n* = 3 biologically independent samples. Error bars, mean ± standard error of mean (SEM).

Furthermore, formation of byproducts was significantly repressed. In ABKS culture, acetate concentrations were decreased 88.2% and 90.4%, and lactate concentrations were decreased 53.2% and 72.7% during the production of PG and 3HP, respectively. The trade-off in productions of target chemicals and byproducts was highlighted in Fig. 2. The correlation of five genetic modifications with the accumulation of lactate and acetate and PG and 3HP production was analyzed (Fig. 2a). Five genetic modifications could inhibit the accumulation of acetate and lactate and promote PG and 3HP production in different degrees, except Δ*iclR* impaired 3HP synthesis. This may explain why the quadruple mutant ABKS (Δ*arcA* Δ*ackA* P_T7_-*csrB* p*acs*) represented the best performance for the productions of PG and 3HP among all the mutants.

**Fig. 2.**
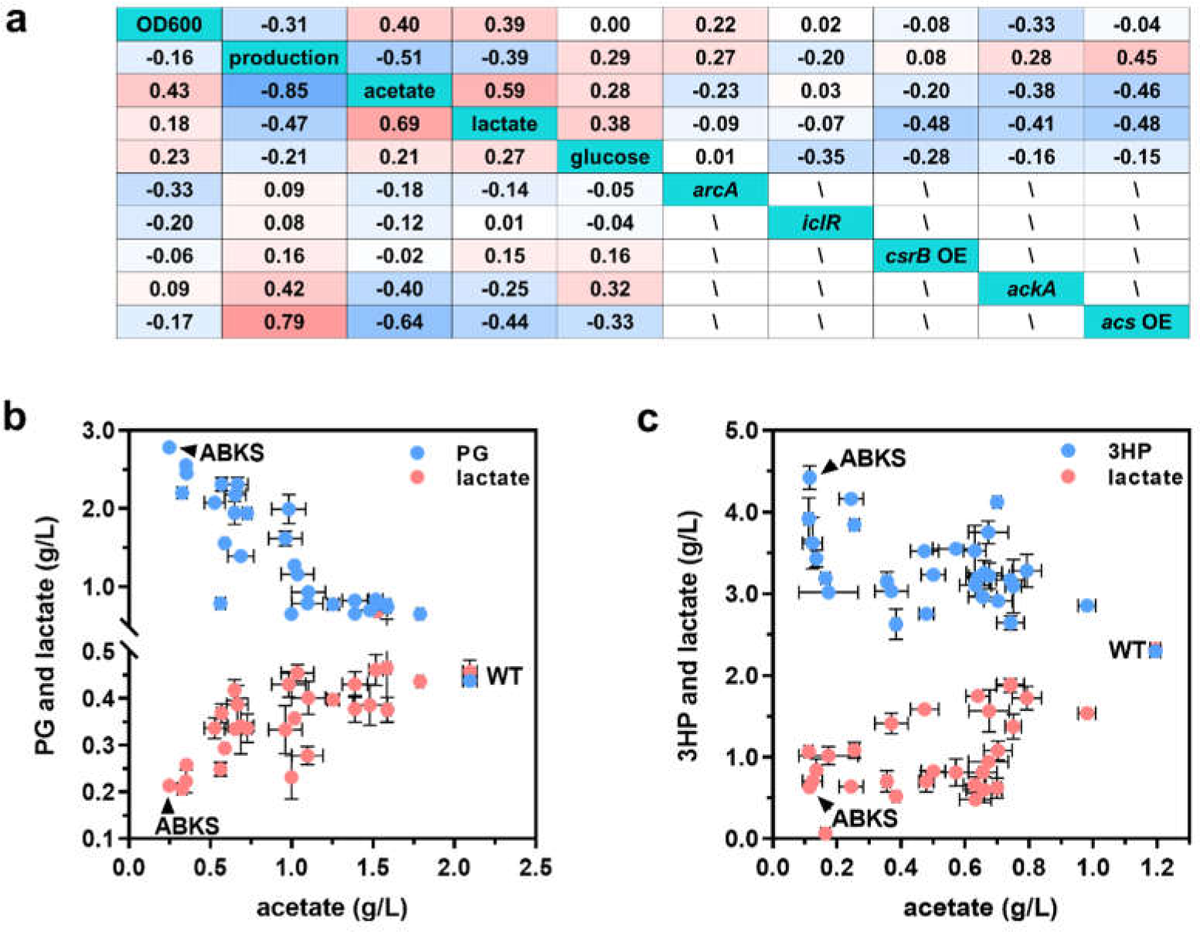
Correlation of genetic modification with target chemicals production. (a) Correlation of five genetic modifications with byproducts accumulation and targets production; The top of the table represents the process of 3HP bioproduction, and the bottom of the table represents the process of PG bioproduction. Effects of acetate overflow on the accumulation of lactate and target chemicals PG (b) and 3HP (c). *n* = 3 biologically independent samples. Error bars, mean ± SEM.

In addition, acetate and lactate accumulation in overflow metabolism led to increased glucose consumption, higher OD_600_ and reduced PG and 3HP titer (Fig. 2a). The effects of acetate overflow on the lactate accumulation and target chemicals production were also analyzed in Figure. 2b and 2c. In these figures, the expected trends are displayed clearly: acetate production correlates positively with lactate production, and correlates negatively with target chemicals production, indicating that acetate and lactate are produced synergistically, and competes the carbon flux from glucose with production of target chemicals. In conclusion, our genetic modifications significantly improved the bioconversion from glucose to desired products, and the strain ABKS with the best performance were selected for further characterization.

### Growth profiles of *E. coli* BL21(DE3) wild-type and ABKS strains

To figure out differences between *E. coli* BL21(DE3) wild-type and ABKS strain, characterizations were carried out using WT-PG (BL21(DE3) strain carrying PG pathway) and ABKS-PG (ABKS strain carrying PG pathway). As reported previously, overflow metabolism will take place when the cells surpass a threshold value of specific glucose consumption rate that is directly related to the specific growth rate of *E. coli* (*12*). When growing in minimal medium with glucose as sole carbon source, WT-PG and ABKS-PG strains both presented a classic S-shaped growth curve (Fig. 3a). The maximum specific growth rate of ABKS-PG strain was a little bit higher than that of WT-PG strain, whereas the maximum specific glucose consumption rate of WT-PG strain was 1.62-time higher than that of ABKS strain (Fig. 3b), suggesting the decoupling of glucose consumption rate and growth rate in ABKS-PG strain. It is believed that this lower specific glucose consumption rate assures ABKS-PG strain of no excretion of byproduct acids at high growth rate.

**Fig. 3.**
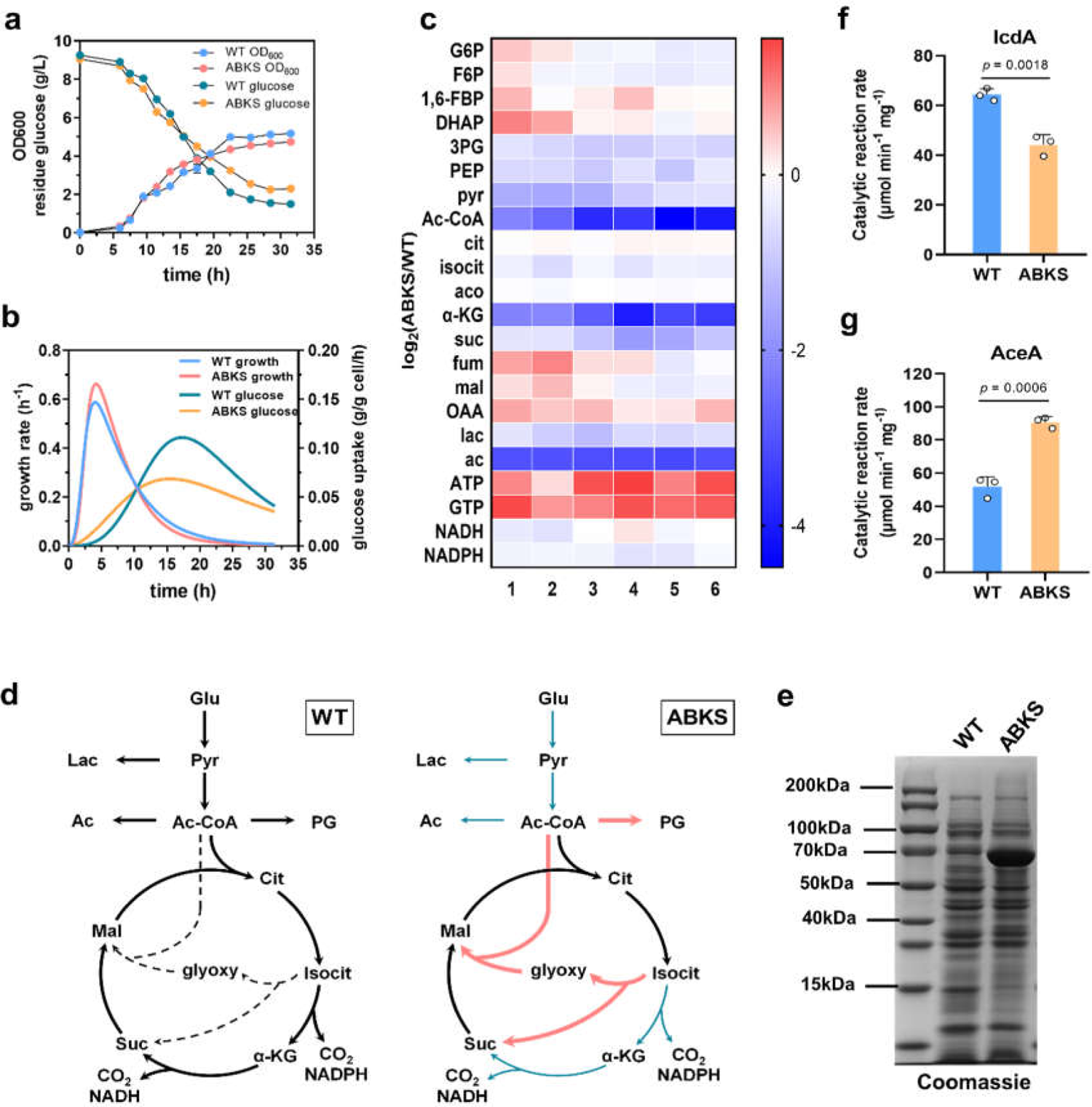
Growth profile and targeted metabolome analysis of WT-PG and ABKS-PG strains. (**a**) Cell density and glucose consumption of WT-PG and ABKS-PG strains. These two strains were inoculated into shaking-flask containing minimal medium with glucose as sole carbon source, and grown at 37 °C (*n* = 3 biologically independent samples). (**b**) Specific growth rate and specific glucose consumption rate of WT-PG and ABKS-PG strains were shown (*n* = 3 biologically independent samples). (**c**) Cluster analysis of intracellular metabolites in WT-PG and ABKS-PG strains of 6 biologically independent samples. (**d**) Proposed reprogrammed central carbon metabolism in WT-PG and ABKS-PG strains. Red line, higher metabolic flux in ABKS-PG than in WT-PG; green line, lower metabolic flux; dashed line, inactive glyoxylate shunt in WT-PG strain. Glu, glucose; G6P, glucose 6-phosphate; F6P, fructose 6-phosphate; 1,6-FBP, fructose 1,6-bisphosphate; DHAP, dihydroxyacetone phosphate; 3PG, glycerate 3-phophote; PEP, phospho*enol*pyruvate; pyr, pyruvate; Ac-CoA, acetyl-CoA; cit, citrate; isocit, isocitrate; aco, *cis*-aconitate; α-KG, α-ketoglutarate; suc, succinate; fum, fumarate; mal, malate; OAA, oxaloacetate; glyoxy, glyoxylate; lac, lactate; ac, acetate. Error bars, mean ± SEM. (e) The whole-cell lysates were normalized to protein concentration, separated by SDS-PAGE, and detected by Coomassie blue stain. Results of the isocitrate dehydrogenase assay (f) and isocitrate lyase assay (g) on crude cell free extracts. (*n* = 3 biologically independent samples). Error bars, mean ± SEM.

### Remodeling of central carbon metabolism in ABKS strain

The targeted metabolome analysis was performed using WT-PG and ABKS-PG strains, and relative amounts of main metabolites in central carbon metabolism and by-products was determined by LC-MS/MS. Consistent with results in Fig. 1a, less acetate and lactate were detected in the ABKS-PG cells (Fig. 3c and Supplementary Fig. 2). The intracellular concentrations of acetyl-CoA and several glycolysis intermediates including glycerate 3-phosphate, phospho*enol*pyruvate (PEP) and pyruvate were significantly lower in ABKS-PG strain than in WT-PG strain. For metabolites in TCA cycle, these two strains presented similar intracellular levels except that the α-ketoglutarate concentration in ABKS-PG strain decreased 85.7 ± 6.9% compared with that in WT-PG strain, indicating that there is alternative route converting isocitrate into succinate. It is speculated that this complementary pathway should be the glyoxylate shunt, although it is normally inactive in *E. coli* with the presence of excess glucose (*10*). In addition, the ABKS strain presented higher intracellular level of ATP and GTP than the wild-type strain.

The metabolome analysis indicated the remodeling of central carbon metabolism in ABKS-PG strain as shown in Fig. 3d. In order to test our hypothesis, the enzyme-catalyzed reaction rates at the isocitrate branching of these two strains were evaluated using crude cell free extracts. The crude cell free extracts were normalized to protein concentration (Fig. 3e). The reaction rate from isocitrate to α-ketoglutarate catalyzed by isocitrate dehydrogenase IcdA was decreased from 64.31 μmol min^-1^ mg^-1^ in WT-PG strain to 44.17 μ mol min^-1^ mg^-1^ in ABKS-PG strain (Fig. 3f). Whereas, the reaction rate from isocitrate to glyoxylate catalyzed by isocitrate lyase AceA in ABKS-PG strain was improved by 1.78 folds as compared to that of WT-PG strain (Fig. 3g). The reaction rate evaluation at the isocitrate branching through an in vitro isocitrate dehydrogenase and lyase assay on crude cell free extracts further illustrated that the metabolic flux may be redistributed from TCA cycle to glyoxylate shunt at the isocitrate branching in ABKS-PG strain.

In conclusion, the glucose uptake and metabolism through glycolysis pathway were down-regulated, leading to decreased intracellular concentrations of pyruvate and acetyl-CoA. Then, main catabolic route of acetyl-CoA switched from TCA cycle to the glyoxylate shunt, shortcutting two decarboxylating steps of TCA cycle and decreasing the CO_2_ release and NAD(P)H production. Furthermore, the glyoxylate shunt is regarded as a main route of acetate assimilation, and its activation would stimulate the use of two-carbon substrates for net synthesis of biomass and target product. In short, this remodeling of *E. coli* central carbon metabolism will avoid excessive intracellular accumulation of pyruvate, acetyl-CoA and reducing equivalents, reduce the overflow metabolism and carbon loss, and improve the production and yield of target chemicals from glucose.

### Transcriptome analysis conforms the carbon metabolism reprograming

To reveal the mechanisms of carbon metabolism reprograming, transcriptome sequencing and annotation were conducted to clarify the differences of gene expression between WT-PG strain and ABKS-PG strain. The raw data has been showed in NCBI Sequence Read Archive (SRA) (Accession number: PRJNA949378). Firstly, hierarchical clustering analysis of the differentially expressed genes (DEGs) was used to determine the gene expression patterns (Supplementary Fig. 3a). The gene expression pattern of ABKS-PG strain was significantly different from WT-PG. Totally, 1353 DEGs were determined in these two strains, among that, 633 genes were upregulated and 720 genes were downregulated in the ABKS-PG strain (Supplementary Fig. 3b). In addition, DEGs were annotated by Gene Ontology (GO) and Kyoto Encyclopedia of Genes and Genomes (KEGG) pathway. According to the corrected p-Value (padj), ten of the most significant changes of GO terms for biological processes (BP), molecular functions (MF), and cellular components (CC) were listed in Supplementary Fig. 4. Results showed that GO terms in BP were changed more obviously than those in CC and MF. Among them, the -log10(padj) of organic acid, carboxylic acid and monocarboxylic acid biosynthetic process in BP was the highest, indicating the most significant difference. Whereas, excepted oxidoreductase activity, acting on the aldehyde or oxo group of donors, -log10(padj) of others GO terms in MF was below 2.20E-05. The central metabolic pathways exhibited significant differences between ABKS-PG strain and WT-PG strain, including PTS, glycolysis/gluconeogenesis, pentose phosphate, glyoxylate shunt and byproducts synthetic pathway according to KEGG enrichment analysis (Fig. 4a and 4b). The GO and KEGG enrichment analysis demonstrated that the gene expression of organic acid biosynthetic process and central metabolism pathways in ABKS-PG strain was dramatically changed as compared to WT-PG strain.

**Fig. 4.**
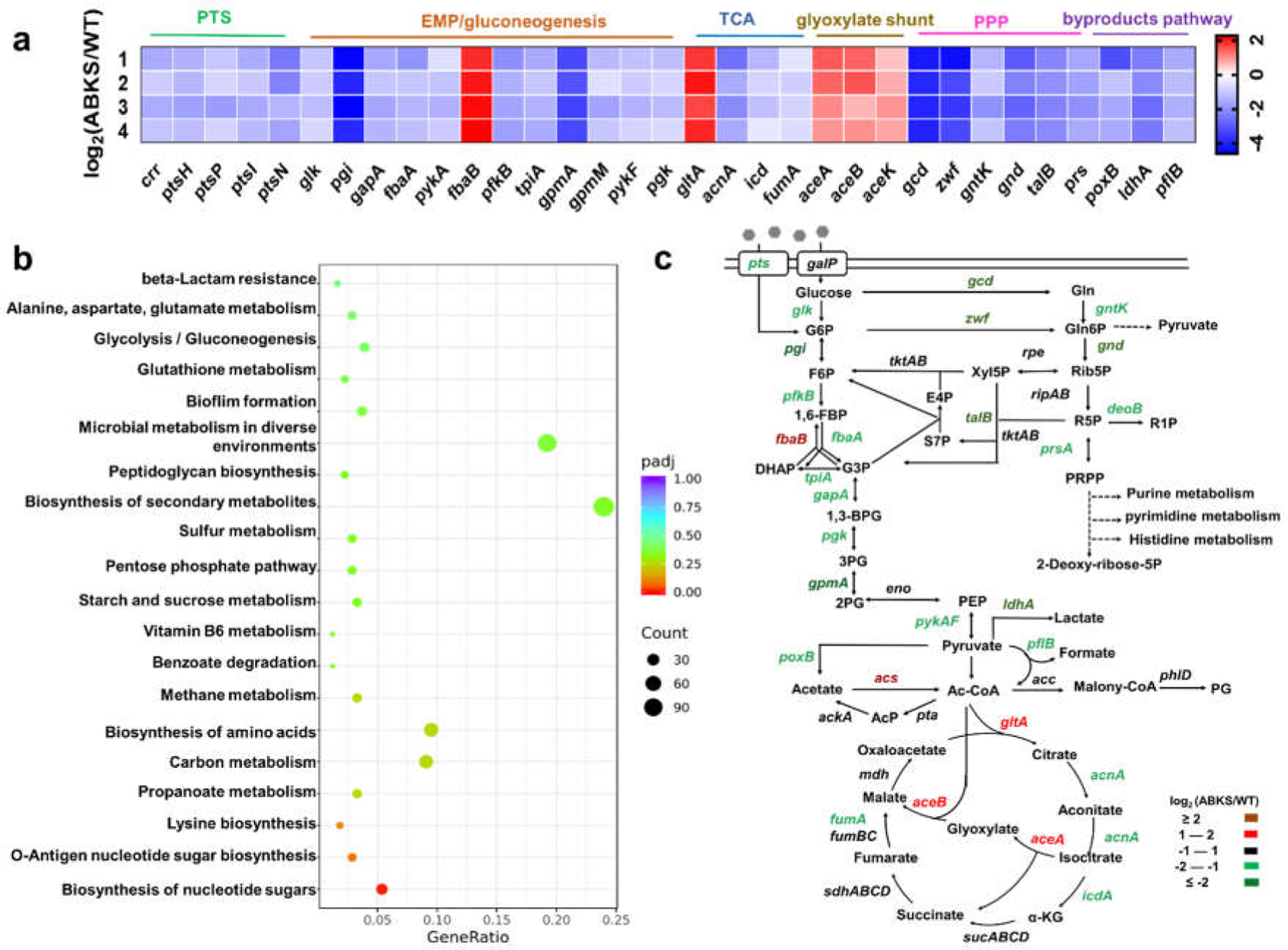
Regulation of central carbon metabolism on gene transcription level in the WT-PG and ABKS-PG strains of 4 biologically independent samples. (a) Cluster analysis related to DEGs from PTS, the central carbon metabolism and by-products synthetic pathways. (b) KEGG enrichment of metabolic genes in DEGs. (c) Sketch of DEGs in central metabolic pathways. G6P, glucose 6-phosphate; F6P, fructose 6-phosphate; 1,6-FBP, fructose 1,6-biphosphate; DHAP, dihydroxyacetone phosphate; G3P, glyceraldehyde 3-phosphate; 1,3-BPG, glycerate 1,3-biphosphate; 3PG, glycerate 3-phophote; 2PG, glycerate 2-phosphate; PEP, phospho*enol*pyruvate; Ac-CoA, acetyl-CoA; AcP, acetyl phosphate; α-KG, α-ketoglutarate; Gln, gluconate; Gln6P, gluconate 6-phosphate; Rib5P, ribulose 5-phosphate; R5P, ribose 5-phosphate; R1P, ribose 1-phosphate; Xyl5P, xylulose 5-phosphate; E4P, erythrose 4-phosphate; S7P, sedoheptulose 7-phosphate; PRPP, 5-phosphoribosyl-1-pyrophosphate.

PTS catalyzes the phosphorylation and transportation of a large number of sugar substrates, and is composed of two common cytoplasmic proteins EI (Enzyme I, encoded by *ptsI* gene) and HPr (encoded by *ptsH* gene), and a carbohydrate-specific complex EII (Enzyme II) (*43*). For glucose, this specific EII complex is encoded by the genes *crr* and *ptsG*. In the ABKS-PG strain, the expression level of *ptsI*, *ptsH* and *crr* genes were downregulated more than 2.5 folds (Fig. 4c and Supplementary Table 2). Furthermore, transcription of genes encoding PTS components for other sugars were also repressed significantly, including *ptsN* and *ptsP* genes.

With function of PTS, glucose is transported into *E. coli* cells and converted into glucose-6-phosphate (G6P), which is a branching-point of glycolysis and pentose phosphate pathway. G6P can be converted into fructose-6-phosphate by phosphoglucose isomerase (encoded by *pgi* gene) then further entering glycolysis pathway, and transcription of *pgi* gene showed the most significant decrease among genes in central metabolism pathways. G6P dehydrogenase Zwf catalyzes the first reaction of pentose phosphate pathway to convert G6P into gluconate-6-phosphate, and the mRNA level of *zwf* gene in ABKS-PG strain only accounted for 8.32% of that in WT-PG strain. Besides PTS, glucose can also be transferred into cytoplasm by galactose permease GalP without phosphorylation (*44*), and then is catalyzed to G6P by glucokinase Glk or to gluconate by glucose dehydrogenase Gcd. In the ABKS-PG strain, both *glk* and *gcd* genes presented a relatively lower mRNA abundance when compared with the WT-PG strain. In addition, genetic modifications of ABKS strain also significantly decreased the expression level of some other genes in glycolysis and pentose phosphate pathways. All these results suggested that expression of enzymes involved in PTS and glucose catabolism was repressed in the ABKS-PG strain, which will reduce the glucose uptake and consumption rate directly. Different from the downregulation of most central metabolism related genes, the expression level of two genes involved in glycolate bypass, isocitrate lyase gene *aceA* and malate synthase gene *aceB*, increased notably in the ABKS-PG strain. As discussed above, activation of the glyoxylate shunt should greatly contribute to lowering the acetate overflow and reducing the carbon loss. Deserved to be mentioned, the synthetic pathways of byproducts, such as acetate, lactate, and formate, were all downregulated in the ABKS-PG strain. Pyruvate oxidase PoxB is the main pathway for acetate production in stationary phase (*45*), and its transcription was severely repressed in the ABKS-PG strain. Furthermore, the genes *pflB* and *ldhA* involved in formation of formate and lactate from pyruvate were downregulated by 2.56 and 5.26 folds, respectively. Coupling with the facts that acetate assimilation enzyme ACS was overexpressed and the *ackA* gene was deleted, these changes in ABKS-PG strain will be benefit for redistributing the overflow carbon flux to our desired pathways.

In summary, transcriptome analysis revealed the downregulation of glucose PTS, glycolysis pathway and TCA cycle and activation of the glyoxylate shunt in the ABKS strain, and provided direct evidence for the reprograming of central carbon metabolism

### Protein acetylation contributes to the carbon metabolism reprograming

Protein acetylation is a conserved posttranslational modification (PTM) in bacteria, and most of the central metabolism enzymes are acetylated, affecting enzyme activity and metabolic flux distribution (*46*). In *E. coli*, protein acetylation occurs via enzymatic and nonenzymatic mechanisms: acetyl group transfer to lysine from acetyl-CoA catalyzed by acetyltransferase PatZ, or from acetyl phosphate (AcP) directly, and a fraction of acetylations are reversed by deacetylase CobB (Fig. 5a). Both intracellular supply of acetyl group donors and expression level of related enzymes affect the protein acetylation. So, protein acetylation and central metabolism affect each other, forming a complex regulatory network, and it was tested whether the protein acetylation status were changed and gave feedback on central metabolism in the ABKS strain. The intracellular concentrations of acetyl-CoA and AcP were both reduced significantly (Fig. 3c and 5b), the mRNA level of acetyltransferase gene *patZ* decreased and the mRNA level of deacetylase gene *cobB* increased remarkably in the AKBS strain (Fig. 5c). These results were entirely consistent with decreased universal protein acetylation level in the ABKS strain (Fig. 5d).

**Fig. 5.**
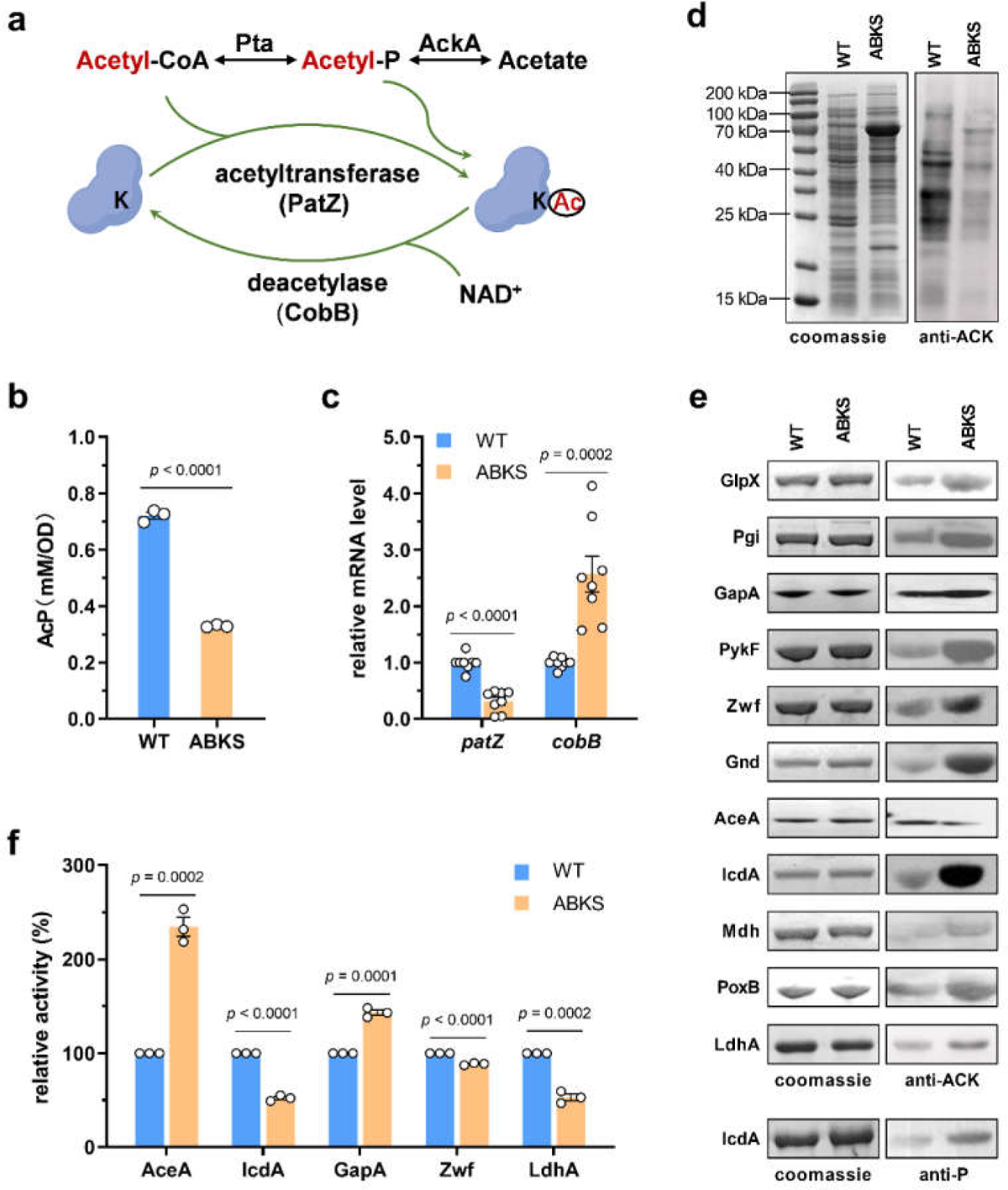
Regulation of protein acetylation and specific activities of some enzymes involved in central metabolism of the WT-PG and ABKS-PG strains. (**a**) Schematics of protein acetylation via enzymatic and nonenzymatic mechanisms in *E. coli*. (**b**) The intracellular acetyl phosphate concentration of the WT and ABKS strains (*n* = 3 biologically independent samples). (**c**) Relative mRNA level of lysine acetyltransferase gene *patZ* and deacetylase gene *cobB* determined by qRT-PCR in the WT and ABKS strains (*n* = 4 biologically independent samples with two technical repeats). (**d**) Anti-acetyllysine immunoblot analyses of whole cell protein of the WT and ABKS strains. The whole-cell lysates were normalized to protein concentration, separated by SDS-PAGE, and detected by Coomassie blue stain and anti-actyllysine (anti-AcK) immunoblot. (**e**) Posttranslational modifications of some metabolic enzymes. The enzyme genes were cloned onto plasmid vector, expressed and purified in the WT and ABKS strain respectively. The same amount of purified protein was applied onto SDS-PAGE, detected by Coomassie blue strain, anti-acetyllysine and anti-phosphoserine (anti-P) immunoblot. (**f**) Specific enzyme activities of some metabolic enzymes purified from the WT and ABKS strains (*n* = 3 biologically independent samples). Error bars, mean ± SEM. **d** and **e** are representative results from two independent experiments.

However, when some enzymes involved in central metabolism were expressed in the WT and ABKS strains, most proteins presented enhanced acetylation status except that the acetylation of isocitrate lyase AceA decreased (Fig. 5e), indicating that the acetylation of metabolic enzymes were regulated via some unclear complex mechanism, not simply determined by the donor supply and known (de)acetylation enzymes. Furthermore, the acetylation modification has different effects on the activities of various enzymes. It was reported that the specific activities of glyceraldehyde 3-phosphate dehydrogenase GapA and malate dehydrogenase Mdh were positively correlated with their acetylation level (*47, 48*), and the activities of isocitrate lyase AceA was negatively correlated with its acetylation (*47*). These were in the line with our results here, and it was also discovered that the specific activities of G6P dehydrogenase Zwf and lactate dehydrogenase LdhA were impaired by acetylation (Fig. 5f). The regulation of isocitrate dehydrogenase IcdA activity by PTM was more complicated, which is affected by both acetylation and phosphorylation. IcdA kinase AceK can phosphorylate IcdA protein, decreasing IcdA specific activity (*47*), and acetylation of IcdA lowers its activity (*49*). In the ABKS strain, both the phosphorylation and acetylation levels of IcdA protein increased (Fig. 5e), which together led to the reduction of overall IcdA activity by 48% compared with in the wild-type strain (Fig. 5f). This PTM-mediated regulation of metabolic enzymes should help to redistribute the metabolic flux into the glyoxylate bypass in the ABKS strain. In summary, the synergistic effect of multi-layer regulation in gene transcription and protein modification ensured the central metabolism reprogramming in the ABKS strain.

### Isocitrate is a key regulatory node

The above results indicated that isocitrate is a key regulatory node for carbon flux redistribution and improved production of target chemicals. To test this hypothesis, an *E. coli* BL21(DE3) mutant strain with deletion of *icdA* gene and overexpression of glyoxylate bypass genes was constructed, however *icdA* mutation severely impaired growth of the engineered strain, instead an *aceB and aceA*-overexpressed strain (AceBA) was used. In addition, isocitrate dehydrogenase kinase/phosphatase, encoded by *aceK* gene, regulates IcdA activity by reversible phosphorylation (*50*). A previous study has isolated the mutant of AceK6(D477N), which selectively lost phosphatase and retained kinase activity (*51*). Phosphorylation of IcdA by AceK kinase activity can decrease its specific activity (*47*). To force carbon flux through the glyoxylate bypass from TCA cycle, *aceA* and *aceK6*-overexpressed strain (AceAK6) was also constructed.

After introduction of PG biosynthetic pathway, the cell growth, glucose consumption and PG production were compared between the AceBA, AceAK6 and wild type. The final OD_600_ of AceBA strain was higher than those of AceAK6 strain and wild type, and the glucose consumption of AceBA and AceAK6 strain was lower as compared to that of the wild type (Fig. 6a). PG productions of AceBA and AceAK6 strain were 0.84 ± 0.047 g/L and 0.77 ± 0.044 g/L respectively, 1.96-time and 1.76-time higher than that of the wild type. The acetate accumulations of AceBA and AceAK6 strain accounted for 50.3% and 38.1% of the WT-PG strain, and lactate accumulations showed no significant differences between these strains. Similarly, the PG yields from glucose of AceBA and AceAK6 strain were increased by 2.51 and 2.27 times than that of wild strain (Fig. 6b). The results revealed that channeling isocitrate into glyoxylate bypass directly did increase the production and yield of target chemical and decrease acetate overflow.

**Fig. 6.**
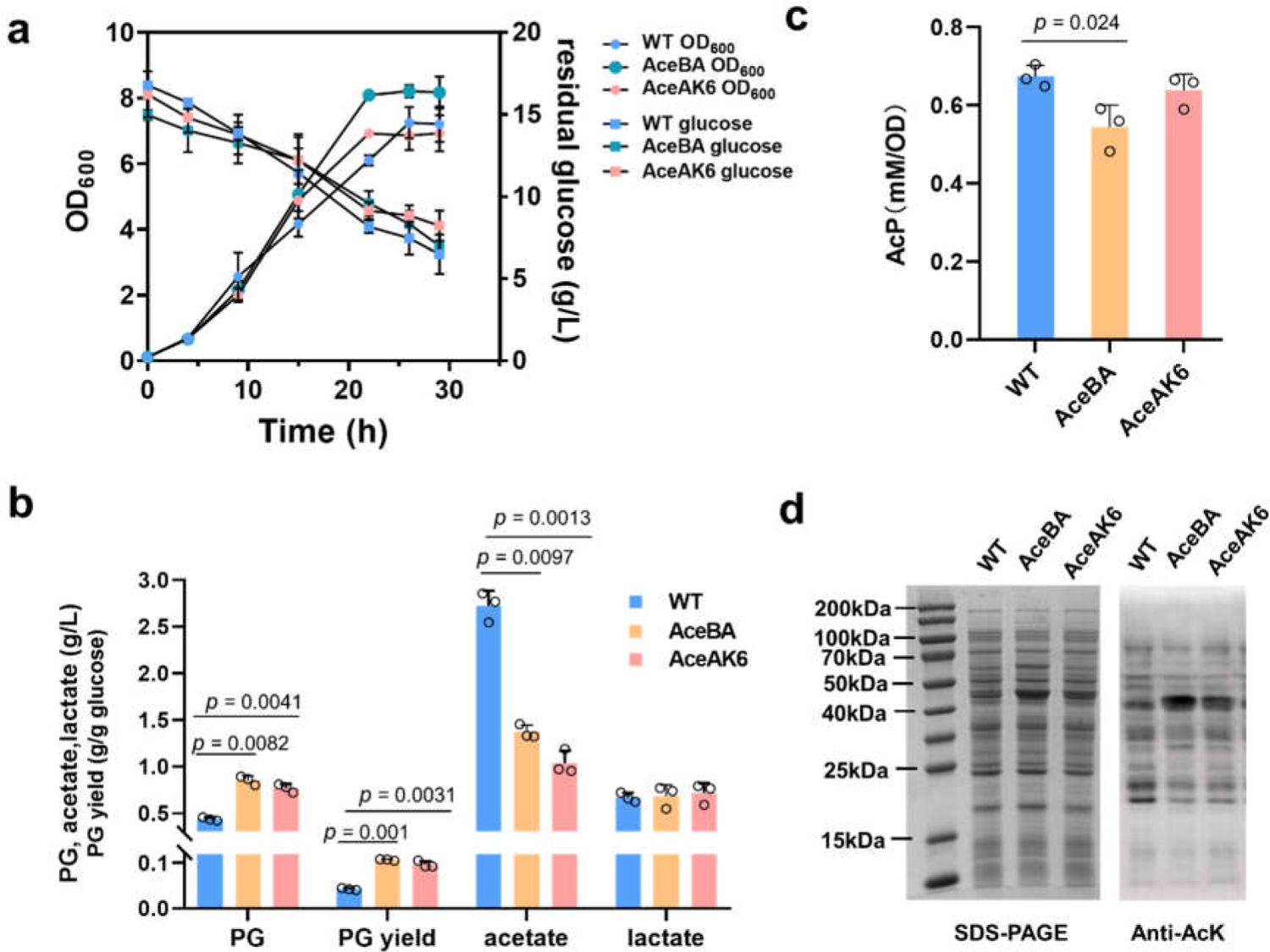
Growth profile and targeted chemical analysis of WT-PG, AceBA-PG and AceAK6-PG strains. (a) Cell density and glucose consumption of the strains. (b) Production of phloroglucinol (PG), acetate and lactate by the strains. (**c**) The intracellular acetyl phosphate concentration of the strains (*n* = 3 biologically independent samples), Error bars, mean ± SEM. (**d**) Anti-acetyllysine immunoblot analyses of whole cell protein of the strains. The whole-cell lysates were normalized to protein concentration.

Protein acetylation plays important roles in enzyme activity and metabolic flux distribution (*46*). We next compared the intracellular AcP concentration and whole protein acetylation level of AceBA-PG and AceAK6 - PG strains with WT-PG. The intracellular AcP concentration of AceBA-PG was slightly lower than that of WT-PG, and there was no significant difference between AceAK6 and WT-PG (Fig. 6c). The whole protein acetylation level of AceBA-PG and AceAK6 - PG was consistent with the changes of intracellular AcP concentration (Fig. 6d). However, both the intracellular concentration of AcP and whole protein acetylation level in ABKS-PG were significantly decreased than those of WT-PG (Fig. 5b and 5d). It could be the main reason why the production and yield of PG in AceBA and AceAK6 strain were lower than that of the ABKS strain. In conclusion, our results have been well demonstrated that isocitrate is a key regulatory node for alleviating the overflow metabolism, reducing carbon loss and improving the production of target chemicals. However, regulating that key node only by overexpressing the genes of glyoxylate bypass at transcriptional level was not enough. Redistributing the metabolic flow at isocitrate node through global and multi-level regulation is able to achieve desirable results.

### The universality of chassis in production of other chemical and protein

To test the universality of chassis, and whether the ABKS strain can improve production of other chemicals, a glycolate biosynthetic pathway from xylose was introduced into BL21(DE3) wild-type and ABKS strain. Besides the endogenous xylose catabolic pathway, these strains can convert xylose into 2-keto-3-deoxy-D-xylonate, followed by an aldolytic cleavage to produce glycolaldehyde and pyruvate. Then glycolaldehyde is oxidized to glycolate by the catalysis of aldehyde dehydrogenase, and pyruvate could be converted into acetyl-CoA and enter TCA cycle (Fig. 7a). When grown in shaking-flask using xylose as sole carbon source, wild-type and ABKS strain produced 2.95 ± 0.07 g/L and 4.44 ± 0.20 g/L glycolate respectively, and the acetate accumulation in ABKS culture was eliminated, resulting in that the ABKS strain presented a glycolate yield of 0.48 ± 0.01 g/g xylose, close to the theoretical yield of 0.51 g/g xylose and 1.6-time higher than that of wild-type strain (Fig. 7b). Moreover, the ABKS strain also accumulated 2.09 ± 0.21 g/L pyruvate, and presented a slightly lower level of lactate production compared with wild-type strain, indicating the reduced metabolic flux in TCA cycle and repressed lactate production in this strain.

**Fig. 7.**
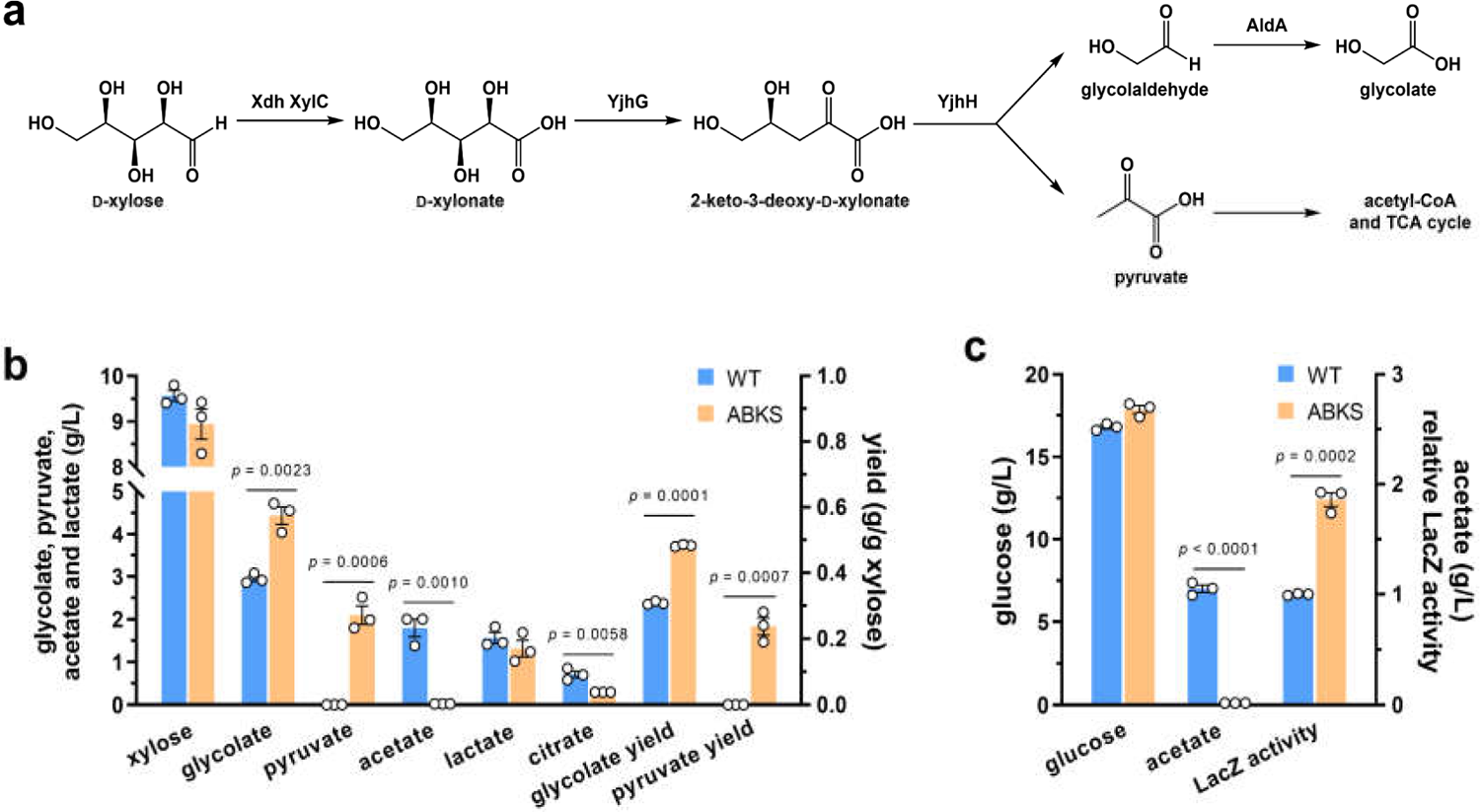
Applications of the ABKS strain in glycolate and β-galactosidase production. (**a**) Glycolate biosynthetic pathway from xylose. Xdh, xylose dehydrogenase; XylC, xlonolactonase; YjhG, xylonate dehydratase; YjhH, 3-keto-3-deoxy-D-xylonate aldolase; AldA, aldehyde dehydrogenase. (**b**) Xylose consumption, glycolate and byproducts production of the BL21(DE3) wild-type and ABKS strain carrying the glycolate pathway (*n* = 3 biologically independent samples). (**c**) Glucose consumption, acetate production and relative LacZ enzyme activity of the BL21(DE3) wild-type and ABKS strain carrying the plasmid pTrcHis2B-*lacZ* (*n* = 3 biologically independent samples). Error bars, mean ± SEM.

In addition to platform chemicals, recombinant proteins for industrial and medical uses are also produced using *E. coli* host. To test whether the ABKS strain is suitable for protein production, β-galactosidase gene *lacZ* was cloned and overexpressed in wild-type and ABKS strain. As show in Fig. 7c, the ABKS strain showed a 1.9-time higher LacZ activity than BL21(DE3) wild-type strain and didn’t accumulate acetate in its culture. All these results suggested that the ABKS strain can effectively reduce the production of byproducts and improve net synthesis of target products, either chemicals or proteins, even when using carbon sources other than glucose.

## Discussion

To convert a living microorganism into a truly productive biofactory, besides optimization of biosynthetic pathway of target product, the endogenous metabolic network of the host, especially its central metabolism, also needs to be tuned to support the bioproduction process. *E. coli* is the host of choice to produce a wide variety of chemicals and proteins. However, the *E. coli* biofactory was usually constructed through optimizing the biosynthetic pathway of target chemical, producing an engineered strain with limited application. It is possible to generate a universal chassis by solving a common obstacle in the endogenous metabolic network, that can draw increasing attention in microbial synthesis.

Overflow metabolism, namely accumulation of acetate and lactate from glucose even in the presence of oxygen, is one of the major drawbacks limiting *E. coli*’s productivity and atom economy. In this study, an *E. coli* chassis of ABKS strain with effectively suppressed overflow metabolism was constructed, presenting remodeled central carbon metabolism and highly improved production and yield of target chemicals and proteins. Compared with previous studies, this strain is expected to have some advantages. Firstly, the ABKS strain has a very wide range of applications, that can be used as a universal chassis for the bioproduction of multiple target chemicals and proteins from different carbon sources. Secondly, the chassis redistributed the metabolic flux from TCA cycle to glyoxylate shunt at the isocitrate branching point. This metabolic reconfiguration bypassed two steps of natural carbon release in TCA cycle, broke the restriction of the original metabolic network and improved the carbon atom economy. Thirdly, genetic modifications in the ABKS strain didn’t cause any unfavorable effect that was usually reported in previous study. For example, deletion of the acetate kinase gene *ackA* or lactate dehydrogenase gene *ldhA* could repress overflow metabolism to some extent, but introduced accumulation of pyruvate in fermentation broth (*25, 26*), whereas our strain didn’t produce any other byproducts. Furthermore, knocking out the genes encoding PTS could slow down the glucose uptake and decrease the acetate accumulation effectively, and these operations led to significantly reduced growth rate of the engineered strain (*28–30*). In opposition, the decoupling of glucose consumption rate and growth rate was observed in the ABKS strain, resulting in a similar growth rate and a remarkably reduced glucose consumption rate compared with the wild-type strain.

Our results proved that holistic engineering of host central metabolism is essential for enhanced bioproduction. There are four genes genetically modified in the ABKS strain, encoding the acetate producing and assimilating enzymes AckA and ACS, and global regulators ArcA and CsrB. Deletion of *ackA* gene and overexpression of *acs* gene should lead to a change of the intracellular level of acetyl-CoA and AcP, which was observed in the ABKS strain. Acetyl-CoA and AcP are both central metabolites and secondary messengers (*52, 53*). Specifically, acetyl-CoA is a key node in metabolism because of its intersection with many metabolic pathways, and influences the activity of multiple enzymes in an allosteric manner (*54*) and protein acetylation in enzymatic mechanism (*46*). Moreover, AcP is not only a critical determinant of nonenzymatic protein acetylation (*53*), but also sufficient for direct phosphorylation of two-component response regulator (*55*), showing regulatory activity on both gene transcription and protein modification levels. For the global regulator ArcA, its deletion improved production of acetyl-CoA-derived chemicals, repressed overflow metabolism (*31*), and significantly enhanced the mRNA level of TCA cycle gens (*56*).

Overexpression of the small regulatory RNA CsrB led to elevation of metabolite and protein levels in glycolysis and TCA cycle, and improved the production of desired compounds by up to two folds (*41*). The ABKS strain constructed in this study shared some similar but not totally consistent phenotypes with each of above mutants, suggesting that should be synergistic effect of those four mutations. It is necessary to carry out further research to elucidate how these genes interplay in the ABKS strain in the future.

Bacteria metabolizes glucose in two main routes: complete oxidation of glucose to CO_2_ and H_2_O through respiration pathway, and incomplete oxidation to byproducts through glycolysis and fermentation. In overflow metabolism, incomplete glucose oxidation is employed instead of complete oxidation even with the presence of oxygen, although incomplete oxidation leads to a several-fold decrease in ATP produced per molecule of glucose (*57*). Recent studies revealed that ATP generation via respiration is limited by finite cellular proteome resources as a series of enzymes in TCA cycle and electron transport chain need to be synthesized, while only a few enzymes are required in fermentation (*33*). So, fermentation is more efficient in the context of the proteome mass required to generate the same amount of ATP, although not carbon-efficient, allowing cells to achieve higher growth rate with excessive glucose. In this study, the ABKS strain presented similar growth rate and enhanced intracellular ATP level compared with BL21(DE3) wild-type strain, without accumulation of fermentation byproducts acetate and lactate, indicating that the proteome cost of energy biogenesis by respiration decreased in this strain. This could be a consequence of the combination of following facts: repressed transcription level of genes in glycolysis and TCA cycle, increased specific activities of metabolic enzymes caused by changed acetylation level, and the switch of carbon flux from TCA cycle to glyoxylate bypass which employs two enzyme isocitrate lyase AceA and malate synthase AceB to shortcut five reactions catalyzed respectively by isocitrate dehydrogenase IcdA, α-ketoglutarate dehydrogenase complex SucAB-Lpd, succinyl-CoA synthetase SucCD, succinate: quinone oxidoreductase SdhABCD and fumarase FumABC, 13 peptides totally. The decreased proteome requirement of respiration pathway assures ABKS strain of a carbon- and proteome-efficient way for energy biogenesis. Therefore, the holistic engineering of host central metabolism in this study enabled the chassis to fulfil the characteristics desirable for industrial bioproduction. In addition, it was discovered that acetylation of metabolic enzymes greatly contributed to rewiring of *E. coli* central metabolism network. In previous metabolic engineering studies, transcriptomic, metabolomic, and proteomic analytic methods were usually used to reveal the physiological and metabolic differences between strains before and after the genetic modification, but little attention has been paid to the changes in acetylation level of proteins. Actually, approximately 90% of bacterial central metabolism enzymes are acetylated, and about 50% of acetylated proteins are involved in diverse metabolic pathways (*47*). Protein acetylation level is deeply affected by central metabolism, dependent on the intracellular supply of acetyl group donor acetyl-CoA and AcP, and the activities of acetyltransferase and deacetylase are regulated by second messengers cyclic adenosine monophosphate (cAMP) (*58*) and cyclic di-guanosine monophosphate (c-di-GMP) (*59*), respectively. On the other side, acetylation regulates the specific activities of metabolism enzymes, especially enzymes which control the direction of gluconeogenesis versus glycolysis and the branching between TCA cycle and glyoxylate shunt, affecting metabolic flux distribution (*47*).

Moreover, protein acetylation can regulate protein amount by affecting the protein aggregation and stability (*60, 61*), and modulate the DNA-centered processes such as DNA replication, repair, and recombination by modifying DNA-binding proteins or related enzymes (*62, 63*). Our previous studies have been established a paradigm for lactate synthesis regulation by protein acetylation (*64, 65*). Altogether, protein acetylation and central metabolism affect each other, forming a regulatory network with high complexity. So, protein acetylation is believed to be a powerful tool for metabolic regulation, and should be paid more attention to in following metabolic engineering studies.

Carbon atom economy, first proposed by Barry M. Trost of Stanford University (*66*), is an important criterion for the biosynthesis process, affecting the efficiency and production cost of biotransformation. An ideal reaction is expected to convert 100 percent of the atoms from the raw material into products, achieving “zero waste emission”. However, CO_2_ release is inevitable in the natural metabolic pathways of microorganisms, like the reaction of pyruvate to acetyl-CoA in EMP pathway and the two decarboxylating steps of TCA cycle, which decreases the carbon atom economy. To address the natural bottleneck of carbon loss from pyruvate to acetyl-CoA in EMP pathway, researchers proposed a non-oxidative glycolysis (NOG) pathway through mathematical calculation, in which 1 mol glucose was directly converted into 3 mol AcP without CO_2_ emission (*67*). CO_2_ release in TCA cycle has become an urgent problem to be solved.

Our results suggested that isocitrate is an important regulatory node for improving the carbon atom economy and production of target chemicals and proteins. Isocitrate lyase AceA splits isocitrate into glyoxylate and succinate, channeling the carbon flow to glyoxylate shunt, and isocitrate dehydrogenase IcdA catalyzes the decarboxylation of isocitrate to α-ketoglutarate, directing the carbon flow into TCA cycle. In the ABKS strain, the AceA protein was regulated in both gene transcription and protein modification levels. The enhanced transcription level of *aceA* gene and reduced acetylation level of AceA, along with decreased IcdA specific activity caused by PTM, facilitated the metabolic flux redistribution at the isocitrate branching point from TCA cycle to glyoxylate shunt. Furthermore, the importance of isocitrate node was also demonstrated by the fact that the glyoxylate bypass-overexpressing strain produced more PG than wild-type strain. The glyoxylate bypass shortcuts two decarboxylating steps of TCA cycle, and stimulates the acetate assimilation, resulting in improved net synthesis of target products and enhanced carbon atom economy. As a result, the proportion of carbon element in ABKS biomass was increased to 51.92%, higher than that of 44.7% in wild strain (Supplementary Fig. 5). Besides *E. coli*, overflow metabolism occurs widely in fast-growing bacteria and fungi (*68*), as well as mammalian stem cells and cancerous cells, also referred to as Crabtree effect in yeast and Warburg effect in cancer cells (*69, 70*). Although the Warburg effect has been well-documented and is of intense interest, its causes or its functions remain unclear. Our results will provide a theoretical basis for the rational design of efficient bioproduction strains and universal chassis, and may shed a light on studies about the Warburg effect in cancer.

In this study, five *E. coli* genes related with central carbon metabolism and acetate metabolism were modified individually or combined randomly, 31 resultant mutants were characterized using biosynthesis of PG and 3HP as proxy, and the ABKS strain with the best performance were selected as the universal chassis for production of chemicals and proteins. In this process, plenty of heavy works of strain construction and testing were involved, which had been expected to be done by some computational tools. Indeed, there are a variety of modeling approaches developed to predict responses of *E. coli* metabolic network to gene perturbation, but several modeling methods presented a very poor overall agreement of predicted and experimentally observed data in previous study (*71*). A possible reason of this low quantitative accuracy is that those modeling approaches all rely on the stoichiometry of metabolic network, hard to tell the effects of genetic modification related to isozymes and global regulators. So, experimental test is still the best and feasible way to determine bacterial physiological response to gene perturbation at nowadays, and it is hoped that computational modeling tools with high accuracy could be developed quickly, relieving microbiologists from the heavy work of strain construction and testing.

## Methods

### Constructions of plasmids and strains

All bacteria strains and plasmids used in this study were listed in Supplementary Table 3, and all primers used were synthesized by Sangon Biotech (Shanghai) Co. Ltd., and listed in Supplementary Table 4. *E. coli* DH5α was used for gene clone, *E. coli* χ7213 strain was used as the host to prepare the suicide vector, and *E. coli* BL21(DE3) was used as the host for production of PG and 3HP. Plasmids are constructed using standard restriction cloning and confirmed by DNA sequencing. P_1_ vir-mediated transduction was used for the knockout of chromosomal genes. The donor strains were purchased from the Keio collection (*72*). The induced overexpression of *csrB* was carried out by inserting the sequence of T_7_-lacO (TAATACGACTCACTATAGGGGAATTGTGAGCGGATAACAATTCC) before *csrB* gene in chromosome using suicide plasmid pRE112-mediated homologous recombination.

### Shake-flask cultivation and analytical methods

Shake-flask experiments were carried out in triplicate series of 250 ml unbaffled Erlenmeyer flasks containing 50 ml fermentation media. The strains were grown overnight at 37 °C in LB broth and 1:50 diluted into 50 ml fermentation media. IPTG was added to 0.1 mM at an OD_600_ of 0.6 and cells were further cultured at 30 °C. 10 mg/L Biotin and 20 mM NaHCO_3_ were also added to the culture for 3HP production. The fermentation and analytical methods of PG (*42*), 3HP (*4*) and glycolate (*73*) were referred to the previous studies.

The whole-cell AcP concentration was measured at 505 nm by forming ferric acetyl-hydroxamate with hydroxamate (*74*). For LacZ activity test, the cells were lysed, mixed with the reaction substrate, O-nitrophenyl-beta-D-galactopyranoside (ONPG), and detected at 420nm (*75*).

### Metabolome analysis

WT-PG and ABKS-PG strains were cultivated in Shake-flasks, after 24h of IPTG induction, *E. coli* cells were collected from 200 µL of culture by centrifugation and the pellets were vortexed in 800 µL precooled methanol/acetonitrile (1:1, v/v). After sonication for 30 min on ice, the mix was stored at −20 °C for 1 h to precipitate proteins. Then the mix was centrifuged for 20 min at 14000 g and 4 °C. After centrifugation, the supernatant was collected, mixed with 200 µL culture supernatant and dried by a vacuum drying system to detect the intracellular and extracellular metabolites simultaneously. The dried samples were dissolved in 100 µL of acetonitrile/ H_2_O (1:1, v/v) and centrifuged for 25 min at 14000 g and 4 °C. Analysis of the targeted metabolites was performed using an LC-MS/MS system consisting of Agilent 1290 Infinity chromatography and AB Sciex QTRAP 5500 mass spectrometer. Ten mM NH_4_COOH and acetonitrile were used as mobile phases A and B, respectively. Metabolites detection were performed by Scientsgene Co., Ltd. (Qingdao, China). The Multiquant software was used to extract chromatographic peak and retention time. The corresponding standard was used to correct the retention time of the target metabolites.

### Transcriptomic analysis

*E. coli* cells were collected after 24h of IPTG induction, total RNA was extracted using the mirVana miRNA Isolation Kit (Ambion) in accordance with the manufacturer’s protocol. RNA integrity was assessed using the RNA Nano 6000 Assay Kit of the Bioanalyzer 2100 system (Agilent Technologies, CA, USA). Sequencing libraries were generated using NEBNext® UltraTM RNA Library Prep Kit for Illumina® (NEB, USA) and index codes were added to attribute sequences to each sample. Sequencing was performed at Beijing Novogene Bioinformatics Technology Co., Ltd. The raw sequence data were filtered by removing reads containing adapter, reads containing poly-N, and low-quality reads. The clean reads were aligned with the genome of *Escherichia coli* BL21(DE3) (accession No. NC_012978.2). Gene expression was quantified as reads per kilobase of coding sequence per million reads (RPKM) algorithm. Differential expression analysis was performed using the DESeq2 R package (1.16.1). Genes with an adjusted P-value <0.05 found by DESeq2 were assigned as differentially expressed.

### Quantitative RT-PCR

Total RNA was extracted using EASYSpin Plus Bacterial RNA quick extract kit (Aidlab Biotechnologies, China) according to the manufacturer’s protocol. RNA concentration was detected by spectrophotometry at 260 nm. Samples were incubated at 42 °C for 10 min with gDNA Eraser to remove the genomic DNA (Takara). PrimeScript RT Reagent Kit was used for RNA reverse transcription and cDNA synthesis. Quantitative RT-PCR was performed with TB Green Premix Ex Taq (Takara) using QuantStudio 1 system (Applied Biosystems). The constitutively transcribed gene *rpoD* was used as a reference control for normalizing the total RNA quantity of different samples. The relative levels of mRNA were calculated using the ΔΔCt method. Data were from three independent biological samples with two technical replicates (*76*).

### Protein expression, purification, and western blotting analysis

*E. coli* cells were grown overnight at 37 °C in LB medium with appropriate antibiotics. The culture was diluted 1:50 into fresh PG-fermentation media with 2% glucose and incubated under the same condition. Protein expression was induced by 0.1 mM IPTG at an OD_600_ of 0.6 and growth was continued for 24 h at 30 °C. Cells were collected by centrifugation and resuspended in phosphate buffer saline buffer (pH 7.5), and disrupted by high pressure. The protein was purified using Ni-NTA His·Bind Column (Novagen). The purified protein was quantified using BCA protein assay kit (Pierce), separated on 12% SDS-PAGE, and transferred to PVDF membrane for 1.5 h at 15V (Millipore). The membrane was incubated with mouse monoclonal anti-acetyl lysine antibody (EasyBio, China) and HRP-conjugated Goat-anti-mouse antibody (EasyBio, China) for acetylation analysis, and mouse monoclonal anti-phosphoserine antibody (sigma) was used for IcdA phosphorylation analysis. Then protein signal was detected using Immobilon Western HRP substrate (Millipore) and Fusion FX6 Imaging System (Vilber, France).

### Enzyme activity Assay

LdhA reaction mixture contained 100 mM phosphate buffer (pH 7.5), 1 mM NADH, 1 mM pyruvate, and 40 nM purified protein. Zwf reaction mixture contained 50mM Tris-Hcl (pH 7.5), 1 mM NADP^+^, 1 mM G6P, 5mM MgCl_2_ and 40 nM purified protein. The specific activity was assayed by measuring the change of absorbance at 340 nm using Spark 20 microplate reader (Tecan). The IcdA enzyme assay was referred to the previous protocol (*49*), and AceA and GapA were assayed as previously described (*47*).

WT-PG and ABKS-PG strains were cultivated in Shake-flasks, after 24h of IPTG induction, *E. coli* cells were collected by centrifugation and resuspended in phosphate buffer saline buffer (pH 7.5), and disrupted by high pressure. The crude cell free extracts were normalized to protein concentration and assayed IcdA and AceA activity.

## Acknowledgements

This study was financially supported by the NSFC (22277068, 32170085, 31961133014), National Key Research and Development Program of China (2021YFC2100503), the Fundamental Research Funds for the Central Universities (G.Z.), Young Scholars Program of Shandong University (M.L.), Distinguished Scholars Program of Shandong University (G.Z.), and Foundation for Innovative Research Groups of State Key Laboratory of Microbial Technology.

## Author contributions

G.Z. designed the experiments. M.L., L.G., M.H., X.F., and Z.Z. performed the experiments. G.Z., M.L., Q.Q. and M.X. analyzed the results. G.Z. and M.L. wrote the manuscript. All authors edited the manuscript before submission.

## References

1. S. Atsumi, T. Hanai, J. C. Liao, Non-fermentative pathways for synthesis of branched-chain higher alcohols as biofuels. Nature 451, 86–89 (2008).

2. S. K. Lee, H. Chou, T. S. Ham, T. S. Lee, J. D. Keasling, Metabolic engineering of microorganisms for biofuels production: from bugs to synthetic biology to fuels. Curr Opin Biotech.19, 556–563 (2008).

3. Q. Wang, P. Yang, C. Liu, Y. Xue, M. Xian, G. Zhao, Biosynthesis of poly(3-hydroxypropionate) from glycerol by recombinant *Escherichia coli*. Bioresour Technol. 131, 548–551 (2013).

4. C. Liu, Y. Ding, R. Zhang, H. Liu, M. Xian, G. Zhao, Functional balance between enzymes in malonyl-CoA pathway for 3-hydroxypropionate biosynthesis. Metab Eng 34, 104–111 (2016).

5. R. Ding, L. Liu, X. Chen, Z. Cui, A. Zhang, D. Ren, L. Zhang, Introduction of two mutations into AroG increases phenylalanine production in *Escherichia coli*. Biotechnol Lett 36, 2103–2108 (2014).

6. C. A. Contador, M. L. Rizk, J. A. Asenjo, J. C. Liao, Ensemble modeling for strain development of L-lysine-producing *Escherichia coli*. Metab Eng 11, 221–233 (2009).

7. S. Zhao, J. A. Jones, D. M. Lachance, N. Bhan, O. Khalidi, S. Venkataraman, Z. Wang, M. A. G. Koffas, Improvement of catechin production in *Escherichia coli* through combinatorial metabolic engineering. Metab Eng 28, 43–53 (2015).

8. X. R. Li, G. Q. Tian, H. J. Shen, J. Z. Liu, Metabolic engineering of *Escherichia coli* to produce zeaxanthin. J Ind Microbiol Biotechnol 42, 627–636 (2015).

9. P. Calero, P. I. Nikel, Chasing bacterial chassis for metabolic engineering: a perspective review from classical to non-traditional microorganisms. Microb Biotechnol 12, 98–124 (2019).

10. V. Bernal, S. Castano-Cerezo, M. Canovas, Acetate metabolism regulation in *Escherichia coli*: carbon overflow, pathogenicity, and beyond. Appl Microbiol Biotechnol 100, 8985–9001 (2016).

11. W. R. Farmer, J. C. Liao, Reduction of aerobic acetate production by *Escherichia coli*. Appl Environ Microbiol 63, 3205–3210 (1997).

12. M. A. Eiteman, E. Altman, Overcoming acetate in *Escherichia coli* recombinant protein fermentations. Trends Biotechnol 24, 530–536 (2006).

13. M. De Mey, G. J. Lequeux, J. J. Beauprez, J. Maertens, E. Van Horen, W. K. Soetaert, P. A. Vanrolleghem, E. J. Vandamme, Comparison of different strategies to reduce acetate formation in *Escherichia coli*. Biotechnol Prog 23, 1053–1063 (2007).

14. P. Landwall, T. Holme, Influence of glucose and dissolved oxygen concentrations on yields of *Escherichia coli* B in dialysis culture. J Gen Microbiol 103, 353–358 (1977).

15. I. E. Gleiser, S. Bauer, Growth of *E. coli* W to high cell concentration by oxygen level linked control of carbon source concentration. Biotechnol Bioeng 23, 1015–1021 (1981).

16. H. G. Hansen, U. Henning, Regulation of pyruvate dehydrogenase activity in *Escherichia coli* K12. Biochim Biophys Acta 122, 355–358 (1966).

17. G. F. Molgat, L. J. Donald, H. W. Duckworth, Chimeric allosteric citrate synthases: construction and properties of citrate synthases containing domains from two different enzymes. Arch Biochem Biophys 298, 238–246 (1992).

18. D. Georgellis, O. Kwon, E. C. Lin, Quinones as the redox signal for the arc two-component system of bacteria. Science 292, 2314–2316 (2001).

19. S. Iuchi, E. C. Lin, *arcA* (*dye*), a global regulatory gene in *Escherichia coli* mediating repression of enzymes in aerobic pathways. Proc Natl Acad Sci U S A 85, 1888–1892 (1988).

20. D. P. Clark, The fermentation pathways of *Escherichia coli*. FEMS Microbiol Rev 63, 223–234 (1989).

21. S. Sun, Y. Ding, M. Liu, M. Xian, G. Zhao, Comparison of glucose, acetate and ethanol as carbon resource for production of poly(3-hydroxybutyrate) and other acetyl-CoA derivatives. Front Bioeng Biotechnol 8, 833 (2020).

22. P. K. Bunch, F. Mat-Jan, N. Lee, D. P. Clark, The *ldhA* gene encoding the fermentative lactate dehydrogenase of *Escherichia coli*. Microbiology 143, 187–195 (1997).

23. S. K. Kim, W. Seong, G. H. Han, D. H. Lee, S. G. Lee, CRISPR interference-guided multiplex repression of endogenous competing pathway genes for redirecting metabolic flux in *Escherichia coli*. Microb Cell Fact 16, 188 (2017).

24. J. H. Kim, C. Wang, H. J. Jang, M. S. Cha, J. E. Park, S. Y. Jo, E. S. Choi, S. W. Kim, Isoprene production by *Escherichia coli* through the exogenous mevalonate pathway with reduced formation of fermentation byproducts. Microb Cell Fact 15, 214 (2016).

25. K. A. Bauer, A. Ben-Bassat, M. Dawson, V. T. de la Puente, J. O. Neway, Improved expression of human interleukin-2 in high-cell-density fermentor cultures of *Escherichia coli* K-12 by a phosphotransacetylase mutant. Appl Environ Microbiol 56, 1296–1302 (1990).

26. T. B. Causey, S. Zhou, K. T. Shanmugam, L. O. Ingram, Engineering the metabolism of *Escherichia coli* W3110 for the conversion of sugar to redox-neutral and oxidized products: homoacetate production. Proc Natl Acad Sci U S A 100, 825–832 (2003).

27. H. Lin, N. M. Castro, G. N. Bennett, K. Y. San, Acetyl-CoA synthetase overexpression in *Escherichia coli* demonstrates more efficient acetate assimilation and lower acetate accumulation: a potential tool in metabolic engineering. Appl Microbiol Biotechnol 71, 870–874 (2006).

28. R. Chen, W. M. Yap, P. W. Postma, J. E. Bailey, Comparative studies of *Escherichia coli* strains using different glucose uptake systems: metabolism and energetics. Biotechnol Bioeng 56, 583–590 (1997).

29. E. Ponce, Effect of growth rate reduction and genetic modifications on acetate accumulation and biomass yields in *Escherichia coli*. J Biosci Bioeng 87, 775–780 (1999).

30. R. R. Gokarn, M. A. Eiteman, E. Altman, Metabolic analysis of *Escherichia coli* in the presence and absence of the carboxylating enzymes phosphoenolpyruvate carboxylase and pyruvate carboxylase. Appl Environ Microbiol 66, 1844–1850 (2000).

31. M. Liu, L. Yao, M. Xian, Y. Ding, H. Liu, G. Zhao, Deletion of *arcA* increased the production of acetyl-CoA-derived chemicals in recombinant *Escherichia coli*. Biotechnol Lett 38, 97–101 (2016).

32. G. N. Vemuri, M. A. Eiteman, E. Altman, Increased recombinant protein production in *Escherichia coli* strains with overexpressed water-forming NADH oxidase and a deleted ArcA regulatory protein. Biotechnol Bioeng 94, 538–542 (2006).

33. M. Basan, S. Hui, H. Okano, Z. Zhang, Y. Shen, J. R. Williamson, T. Hwa, Overflow metabolism in *Escherichia coli* results from efficient proteome allocation. Nature 528, 99–104 (2015).

34. B. Enjalbert, M. Cocaign-Bousquet, J. C. Portais, F. Letisse, Acetate exposure determines the diauxic behavior of *Escherichia coli* during the glucose-acetate transition. J Bacteriol 197, 3173–3181 (2015).

35. B. Schilling, D. Christensen, R. Davis, A. K. Sahu, L. I. Hu, A. Walker-Peddakotla, D. J. Sorensen, B. Zemaitaitis, B. W. Gibson, A. J. Wolfe, Protein acetylation dynamics in response to carbon overflow in *Escherichia coli*. Mol Microbiol 98, 847–863 (2015).

36. A. F. Alvarez, D. Georgellis, In vitro and in vivo analysis of the ArcB/A redox signaling pathway. Methods Enzymol 471, 205–228 (2010).

37. R. Malpica, G. R. Sandoval, C. Rodriguez, B. Franco, D. Georgellis, Signaling by the Arc two-component system provides a link between the redox state of the quinone pool and gene expression. Antioxid Redox Signal 8, 781–795 (2006).

38. G. L. Lorca, A. Ezersky, V. V. Lunin, J. R. Walker, S. Altamentova, E. Evdokimova, M. Vedadi, A. Bochkarev, A. Savchenko, Glyoxylate and pyruvate are antagonistic effectors of the *Escherichia coli* IclR transcriptional regulator. J Biol Chem 282, 16476–16491 (2007).

39. P. Babitzke, T. Romeo, CsrB sRNA family: sequestration of RNA-binding regulatory proteins. Curr Opin Microbiol 10, 156–163 (2007).

40. M. Liu, Y. Ding, H. Chen, Z. Zhao, H. Liu, M. Xian, G. Zhao, Improving the production of acetyl-CoA-derived chemicals in *Escherichia coli* BL21(DE3) through *iclR* and *arcA* deletion. BMC microbiology 17, 10 (2017).

41. A. E. McKee, B. J. Rutherford, D. C. Chivian, E. K. Baidoo, D. Juminaga, D. Kuo, P. I. Benke, J. A. Dietrich, S. M. Ma, A. P. Arkin, C. J. Petzold, P. D. Adams, J. D. Keasling, S. R. Chhabra, Manipulation of the carbon storage regulator system for metabolite remodeling and biofuel production in *Escherichia coli*. Microb Cell Fact 11, 79 (2012).

42. Y. J. Cao, X. L. Jiang, R. B. Zhang, M. Xian, Improved phloroglucinol production by metabolically engineered *Escherichia coli*. Appl Microbiol Biotechnol 91, 1545–1552 (2011).

43. J. Deutscher, C. Francke, P. W. Postma, How phosphotransferase system-related protein phosphorylation regulates carbohydrate metabolism in bacteria. Microbiol Mol Biol Rev 70, 939–1031 (2006).

44. S. S. Pao, I. T. Paulsen, M. H. Saier, Jr., Major facilitator superfamily. Microbiol Mol Biol Rev 62, 1–34 (1998).

45. C. R. Dittrich, G. N. Bennett, K. Y. San, Characterization of the acetate-producing pathways in *Escherichia coli*. Biotechnol Prog 21, 1062–1067 (2005).

46. M. Liu, L. Guo, Y. Fu, M. Huo, Q. Qi, G. Zhao, Bacterial protein acetylation and its role in cellular physiology and metabolic regulation. Biotechnol Adv 53, 107842 (2021).

47. Q. Wang, Y. Zhang, C. Yang, H. Xiong, Y. Lin, J. Yao, H. Li, L. Xie, W. Zhao, Y. Yao, Z. B. Ning, R. Zeng, Y. Xiong, K. L. Guan, S. Zhao, G. P. Zhao, Acetylation of metabolic enzymes coordinates carbon source utilization and metabolic flux. Science 327, 1004–1007 (2010).

48. S. Venkat, C. Gregory, J. Sturges, Q. Gan, C. Fan, Studying the lysine acetylation of malate dehydrogenase. J Mol Biol 429, 1396–1405 (2017).

49. S. Venkat, H. Chen, A. Stahman, D. Hudson, P. McGuire, Q. Gan, C. Fan, Characterizing Lysine Acetylation of Isocitrate Dehydrogenase in Escherichia coli. J Mol Biol 430, 1901–1911 (2018).

50. J. H. Hurley, A. M. Dean, J. L. Sohl, D. E. Koshland, Jr., R. M. Stroud, Regulation of an enzyme by phosphorylation at the active site. Science 249, 1012–1016 (1990).

51. T. P. Ikeda, E. Houtz, D. C. LaPorte, Isocitrate dehydrogenase kinase/phosphatase: identification of mutations which selectively inhibit phosphatase activity. J Bacteriol 174, 1414–1416 (1992).

52. F. Pietrocola, L. Galluzzi, J. M. Bravo-San Pedro, F. Madeo, G. Kroemer, Acetyl coenzyme A: a central metabolite and second messenger. Cell Metab 21, 805–821 (2015).

53. B. T. Weinert, V. Iesmantavicius, S. A. Wagner, C. Schölz, B. Gummesson, P. Beli, T. Nyström, C. Choudhary, Acetyl-phosphate is a critical determinant of lysine acetylation in E. coli. Mol Cell 51, 265–272 (2013).

54. A. Krivoruchko, Y. Zhang, V. Siewers, Y. Chen, J. Nielsen, Microbial acetyl-CoA metabolism and metabolic engineering. Metab Eng 28, 28–42 (2015).

55. A. H. Klein, A. Shulla, S. A. Reimann, D. H. Keating, A. J. Wolfe, The intracellular concentration of acetyl phosphate in *Escherichia coli* is sufficient for direct phosphorylation of two-component response regulators. J Bacteriol 189, 5574–5581 (2007).

56. T. Oshima, H. Aiba, Y. Masuda, S. Kanaya, M. Sugiura, B. L. Wanner, H. Mori, T. Mizuno, Transcriptome analysis of all two-component regulatory system mutants of *Escherichia coli* K-12. Mol Microbiol 46, 281–291 (2002).

57. T. Pfeiffer, S. Schuster, S. Bonhoeffer, Cooperation and competition in the evolution of ATP-producing pathways. Science 292, 504–507 (2001).

58. H. Xu, S. S. Hegde, J. S. Blanchard, Reversible acetylation and inactivation of *mycobacterium tuberculosis* acetyl-CoA synthetase is dependent on cAMP. Biochemistry 50, 5883–5892 (2011).

59. Z. Xu, H. Zhang, X. Zhang, H. Jiang, C. Liu, F. Wu, L. Qian, B. Hao, D. M. Czajkowsky, S. Guo, Z. Xu, L. Bi, S. Wang, H. Li, M. Tan, W. Yan, L. Feng, J. Hou, S. C. Tao, Interplay between the bacterial protein deacetylase CobB and the second messenger c-di-GMP. Embo J 38, e100948 (2019).

60. D. Kuczynska-Wisnik, M. Moruno-Algara, K. Stojowska-Swedrzynska, E. Laskowska, The effect of protein acetylation on the formation and processing of inclusion bodies and endogenous protein aggregates in *Escherichia coli* cells. Microb Cell Fact 15, 189 (2016).

61. Y. Sang, J. Ren, R. Qin, S. Liu, Z. Cui, S. Cheng, X. Liu, J. Lu, J. Tao, Y. F. Yao, Acetylation regulating protein stability and DNA-binding ability of HilD, thus modulating *Salmonella Typhimurium* virulence. J Infect Dis 216, 1018–1026 (2017).

62. S. Ghosh, B. Padmanabhan, C. Anand, V. Nagaraja, Lysine acetylation of the Mycobacterium tuberculosis HU protein modulates its DNA binding and genome organization. Mol Microbiol 100, 577–588 (2016).

63. Q. Zhou, Y. N. Zhou, D. J. Jin, Y. C. Tse-Dinh, Deacetylation of topoisomerase I is an important physiological function of *E. coli* CobB. Nucleic Acids Res 45, 5349–5358 (2017).

64. M. Liu, M. Huo, C. Liu, L. Guo, Y. Ding, Q. Ma, Q. Qi, M. Xian, G. Zhao, Lysine acetylation of Escherichia coli lactate dehydrogenase regulates enzyme activity and lactate synthesis. Front Bioeng Biotech. 10, 966062 (2022).

65. M. Liu, M. Huo, L. Guo, Y. Fu, M. Xian, Q. Qi, W. Liu, G. Zhao, Lysine acetylation decreases enzyme activity and protein level of Escherichia coli lactate dehydrogenase. Eng Microb 2, (2022).

66. B. M. Trost, On inventing reactions for atom economy. Accounts Chem Res 35, 695–705 (2002).

67. I. W. Bogorad, T. S. Lin, J. C. Liao, Synthetic non-oxidative glycolysis enables complete carbon conservation. Nature 502, 693–697 (2013).

68. C. Malina, R. Yu, J. Bjorkeroth, E. J. Kerkhoven, J. Nielsen, Adaptations in metabolism and protein translation give rise to the Crabtree effect in yeast. Proc Natl Acad Sci U S A 118, (2021).

69. M. G. Vander Heiden, L. C. Cantley, C. B. Thompson, Understanding the Warburg Effect: the metabolic requirements of cell proliferation. Science 324, 1029–1033 (2009).

70. M. V. Liberti, J. W. Locasale, The Warburg Effect: how does it benefit cancer cells? Trends Biochem Sci 41, 211–218 (2016).

71. C. P. Long, J. E. Gonzalez, N. R. Sandoval, M. R. Antoniewicz, Characterization of physiological responses to 22 gene knockouts in *Escherichia coli* central carbon metabolism. Metab Eng 37, 102–113 (2016).

72. T. Baba, T. Ara, M. Hasegawa, Y. Takai, Y. Okumura, M. Baba, K. A. Datsenko, M. Tomita, B. L. Wanner, H. Mori, Construction of Escherichia coli K-12 in-frame, single-gene knockout mutants: the Keio collection. Mol Syst Biol 2, 2006.0008 (2006).

73. M. Liu, Y. Ding, M. Xian, G. Zhao, Metabolic engineering of a xylose pathway for biotechnological production of glycolate in *Escherichia coli*. Microb Cell Fact 17, 51 (2018).

74. Q. Wang, J. Xu, Z. Sun, Y. Luan, Y. Li, J. Wang, Q. Liang, Q. Qi, Engineering an *in vivo* EP-bifido pathway in *Escherichia coli* for high-yield acetyl-CoA generation with low CO_2_ emission. Metab Eng 51, 79–87 (2019).

75. S. T. Smale, Beta-galactosidase assay. Cold Spring Harb Protoc 2010, pdb prot5423 (2010).

76. K. J. Livak, T. D. Schmittgen, Analysis of relative gene expression data using real-time quantitative PCR and the 2(-Delta Delta C(T)) Method. Methods (San Diego, Calif.) 25, 402–408 (2001).

