## Supplementary Information for "Remodeling the central metabolism of *Escherichia coli* enables a universal chassis"

**Supplementary Table 1** Productions of phloroglucinol (PG), 3-hydroxypropionate (3HP), acetate and lactate by *E. coli* BL21(DE3) wild-type strain and 29 genetically modified strains. Data from 3 biological independent samples were shown.

| genotype | | | | | PG production | | | | | | | | | | | | 3HP production | | | | | | | | | | | |
|---|---|---|---|---|---|---|---|---|---|---|---|---|---|---|---|---|---|---|---|---|---|---|---|---|---|---|---|---|
| *arcA* | *iclR* | *csrB* | *ackA* | *acs* | PG (g/L) | | | acetate (g/L) | | | lactate (g/L) | | | PG yield (g/g) | | | 3HP (g/L) | | | acetate (g/L) | | | lactate (g/L) | | | 3HP yield (g/g) | | |
| | | | | | 0.42 | 0.44 | 0.45 | 2.09 | 2.17 | 2.02 | 0.42 | 0.50 | 0.45 | 0.05 | 0.05 | 0.06 | 2.20 | 2.33 | 2.36 | 1.16 | 1.21 | 1.22 | 2.39 | 2.22 | 2.36 | 0.16 | 0.17 | 0.15 |
| + | | | | | 0.90 | 0.89 | 1.00 | 1.30 | 1.08 | 0.93 | 0.45 | 0.42 | 0.33 | 0.09 | 0.09 | 0.10 | 3.42 | 3.54 | 3.61 | 0.42 | 0.56 | 0.44 | 1.63 | 1.51 | 1.63 | 0.27 | 0.29 | 0.28 |
| | + | | | | 0.68 | 0.67 | 0.74 | 1.62 | 1.54 | 1.28 | 0.47 | 0.32 | 0.37 | 0.07 | 0.07 | 0.07 | 3.01 | 3.23 | 3.28 | 0.67 | 0.82 | 0.74 | 2.02 | 1.89 | 1.72 | 0.22 | 0.25 | 0.27 |
| | | + | | | 0.75 | 0.77 | 0.79 | 1.59 | 1.54 | 1.63 | 0.69 | 0.38 | 0.32 | 0.07 | 0.07 | 0.08 | 2.88 | 3.53 | 3.44 | 0.88 | 0.73 | 0.77 | 1.83 | 1.90 | 1.44 | 0.18 | 0.23 | 0.24 |
| | | | + | | 0.61 | 0.70 | 0.64 | 1.00 | 0.99 | 1.01 | 0.30 | 0.25 | 0.14 | 0.09 | 0.08 | 0.08 | 2.73 | 2.47 | 2.73 | 0.82 | 0.72 | 0.68 | 1.89 | 1.81 | 1.97 | 0.20 | 0.24 | 0.27 |
| | | | | + | 0.73 | 0.76 | 0.72 | 1.62 | 1.55 | 1.60 | 0.33 | 0.38 | 0.42 | 0.08 | 0.09 | 0.08 | 3.51 | 2.96 | 3.19 | 0.61 | 0.62 | 0.80 | 1.23 | 1.39 | 2.07 | 0.26 | 0.20 | 0.21 |
| + | + | | | | 0.79 | 0.79 | 0.78 | 0.54 | 0.62 | 0.53 | 0.26 | 0.22 | 0.27 | 0.08 | 0.08 | 0.08 | 3.06 | 3.24 | 3.30 | 0.57 | 0.64 | 0.71 | 1.86 | 1.65 | 1.74 | 0.20 | 0.27 | 0.28 |
| + | | + | | | 0.62 | 0.69 | 0.64 | 1.40 | 1.39 | 1.37 | 0.43 | 0.33 | 0.38 | 0.07 | 0.07 | 0.07 | 3.09 | 3.05 | 2.91 | 0.08 | 0.08 | 0.36 | 1.23 | 0.88 | 0.93 | 0.30 | 0.29 | 0.29 |
| + | | | + | | 1.66 | 2.16 | 2.02 | 0.64 | 0.63 | 0.68 | 0.44 | 0.44 | 0.37 | 0.18 | 0.17 | 0.18 | 3.19 | 3.61 | 2.48 | 0.73 | 0.80 | 0.72 | 1.66 | 1.33 | 1.13 | 0.27 | 0.29 | 0.19 |
| + | | | | + | 1.51 | 1.40 | 1.27 | 0.70 | 0.82 | 0.54 | 0.46 | 0.27 | 0.30 | 0.17 | 0.18 | 0.17 | 3.24 | 3.46 | 3.60 | 0.13 | 0.14 | 0.15 | 1.08 | 0.86 | 0.59 | 0.28 | 0.30 | 0.28 |
| | + | + | | | 0.80 | 0.75 | 0.78 | 1.55 | 1.49 | 1.53 | 0.56 | 0.75 | 0.74 | 0.06 | 0.07 | 0.06 | 3.56 | 3.17 | 3.04 | 0.73 | 0.70 | 0.55 | 0.85 | 0.66 | 0.30 | 0.29 | 0.31 | 0.29 |
| | + | | + | | 0.75 | 0.74 | 0.87 | 1.27 | 1.08 | 0.94 | 0.24 | 0.28 | 0.31 | 0.06 | 0.04 | 0.05 | 2.96 | 3.07 | 3.07 | 0.18 | 0.19 | 0.74 | 1.55 | 1.53 | 1.16 | 0.25 | 0.24 | 0.23 |
| | + | | | + | 1.59 | 1.79 | 1.47 | 0.99 | 1.12 | 0.77 | 0.42 | 0.33 | 0.24 | 0.21 | 0.23 | 0.16 | 4.02 | 3.54 | 3.71 | 0.78 | 0.56 | 0.68 | 0.95 | 1.01 | 0.87 | 0.34 | 0.25 | 0.25 |
| | | + | + | | 0.81 | 0.92 | 0.74 | 1.24 | 1.43 | 1.50 | 0.38 | 0.45 | 0.46 | 0.05 | 0.05 | 0.05 | 3.17 | 3.22 | 3.18 | 0.14 | 0.16 | 0.19 | 0.07 | 0.07 | 0.06 | 0.31 | 0.32 | 0.29 |
| | | + | | + | 1.66 | 1.55 | 1.47 | 0.61 | 0.60 | 0.56 | 0.30 | 0.28 | 0.30 | 0.20 | 0.18 | 0.17 | 2.97 | 2.97 | 3.38 | 0.57 | 0.73 | 0.60 | 0.55 | 0.56 | 0.34 | 0.31 | 0.28 | 0.32 |
| | | | + | + | 1.97 | 2.16 | 2.10 | 0.41 | 0.55 | 0.62 | 0.32 | 0.38 | 0.31 | 0.24 | 0.23 | 0.25 | 3.82 | 3.89 | 3.83 | 0.25 | 0.23 | 0.28 | 1.02 | 0.96 | 1.28 | 0.31 | 0.28 | 0.26 |
| + | + | + | | | 0.63 | 0.65 | 0.66 | 1.75 | 1.82 | 1.79 | 0.42 | 0.46 | 0.43 | 0.07 | 0.07 | 0.07 | 2.86 | 2.92 | 2.79 | 0.94 | 1.03 | 0.98 | 1.64 | 1.49 | 1.48 | 0.28 | 0.26 | 0.28 |
| + | + | | + | | 0.80 | 0.77 | 0.75 | 1.26 | 1.19 | 1.32 | 0.40 | 0.41 | 0.39 | 0.11 | 0.10 | 0.09 | 2.93 | 3.05 | 2.90 | 0.57 | 0.67 | 0.72 | 0.80 | 0.81 | 0.83 | 0.27 | 0.27 | 0.27 |
| + | | + | + | | 0.80 | 0.81 | 0.89 | 1.60 | 1.46 | 1.50 | 0.40 | 0.49 | 0.50 | 0.06 | 0.05 | 0.05 | 4.20 | 4.15 | 4.03 | 0.70 | 0.68 | 0.72 | 0.87 | 0.46 | 0.54 | 0.26 | 0.22 | 0.30 |

| | | | | | | | | | | | | | | | | | | | | | | | | | | | | |
|---|---|---|---|---|---|---|---|---|---|---|---|---|---|---|---|---|---|---|---|---|---|---|---|---|---|---|---|---|
| + | | | + | + | 1.67 | 2.02 | 2.30 | 1.17 | 0.98 | 0.80 | 0.38 | 0.45 | 0.46 | 0.16 | 0.18 | 0.16 | 4.14 | 3.33 | 3.11 | 0.57 | 0.69 | 0.64 | 0.86 | 0.54 | 0.59 | 0.30 | 0.28 | 0.24 |
| | + | + | + | | 1.24 | 1.11 | 1.13 | 0.83 | 1.12 | 1.16 | 0.42 | 0.45 | 0.49 | 0.12 | 0.11 | 0.10 | 2.25 | 2.82 | 2.80 | 0.39 | 0.40 | 0.36 | 0.47 | 0.63 | 0.45 | 0.18 | 0.28 | 0.24 |
| | + | + | | + | 2.23 | 2.16 | 2.14 | 0.59 | 0.59 | 0.78 | 0.33 | 0.33 | 0.35 | 0.24 | 0.25 | 0.26 | 3.01 | 2.95 | 2.78 | 0.65 | 0.67 | 0.79 | 1.28 | 0.87 | 1.09 | 0.30 | 0.31 | 0.24 |
| | + | | + | + | 2.17 | 2.49 | 2.26 | 0.60 | 0.60 | 0.80 | 0.36 | 0.47 | 0.33 | 0.24 | 0.25 | 0.24 | 3.66 | 3.47 | 3.52 | 0.71 | 0.55 | 0.46 | 0.62 | 1.15 | 0.67 | 0.31 | 0.34 | 0.35 |
| | | + | + | + | 2.69 | 2.54 | 2.44 | 0.35 | 0.36 | 0.34 | 0.27 | 0.18 | 0.22 | 0.28 | 0.25 | 0.25 | 4.13 | 4.14 | 4.22 | 0.21 | 0.32 | 0.21 | 0.60 | 0.69 | 0.63 | 0.32 | 0.36 | 0.35 |
| + | + | + | + | | 1.26 | 1.29 | 1.27 | 1.04 | 1.00 | 1.02 | 0.37 | 0.35 | 0.35 | 0.13 | 0.12 | 0.12 | 3.20 | 3.22 | 3.29 | 0.56 | 0.52 | 0.43 | 0.81 | 0.89 | 0.78 | 0.30 | 0.28 | 0.27 |
| + | + | + | | + | 1.94 | 1.84 | 2.03 | 0.69 | 0.78 | 0.71 | 0.30 | 0.31 | 0.40 | 0.21 | 0.20 | 0.21 | 3.38 | 3.01 | 3.06 | 0.37 | 0.36 | 0.33 | 0.59 | 0.97 | 0.55 | 0.27 | 0.22 | 0.24 |
| + | + | | + | + | 2.19 | 2.29 | 2.13 | 0.39 | 0.30 | 0.30 | 0.18 | 0.21 | 0.23 | 0.25 | 0.19 | 0.21 | 3.23 | 4.24 | 3.40 | 0.16 | 0.06 | 0.15 | 0.74 | 0.62 | 0.76 | 0.27 | 0.37 | 0.34 |
| + | | + | + | + | 2.69 | 2.79 | 2.88 | 0.25 | 0.25 | 0.24 | 0.22 | 0.21 | 0.21 | 0.31 | 0.28 | 0.30 | 4.20 | 4.70 | 4.38 | 0.11 | 0.13 | 0.10 | 0.66 | 0.69 | 0.56 | 0.36 | 0.39 | 0.35 |
| | + | + | + | + | 2.23 | 2.49 | 2.22 | 0.52 | 0.57 | 0.62 | 0.35 | 0.34 | 0.41 | 0.28 | 0.29 | 0.25 | 2.66 | 2.74 | 2.86 | 0.46 | 0.45 | 0.52 | 0.59 | 0.97 | 0.55 | 0.32 | 0.28 | 0.31 |
| + | + | + | + | + | 2.36 | 2.46 | 2.54 | 0.35 | 0.36 | 0.35 | 0.27 | 0.23 | 0.28 | 0.27 | 0.26 | 0.29 | 4.40 | 3.53 | 3.83 | 0.09 | 0.11 | 0.14 | 1.02 | 1.15 | 1.04 | 0.37 | 0.32 | 0.34 |

**Supplementary Table 2** Genes expressed differently in WT-PG and ABKS-PG strains

| Gene name | Fold change (ABKS vs wt) | *p* value | Description |
|---|---|---|---|
| **Phosphotransferase system** | | | |
| *ptsN* | 0.278 | 3.75E-02 | sugar-specific enzyme IIA component of PTS |
| *ptsH* | 0.374 | 1.81E-04 | phosphotransferase component of PTS system (Hpr) |
| *ptsI* | 0.402 | 2.20E-03 | PEP-protein phosphotransferase of PTS system (enzyme I) |
| *crr* | 0.422 | 6.29E-03 | glucose-specific enzyme IIA component of PTS |
| *ptsP* | 0.483 | 1.40E-03 | PEP-protein phosphotransferase enzyme I |
| **Glycolysis/Gluconeogenesis pathway** | | | |
| *fbaB* | 4.333 | 1.85E-05 | fructose-bisphosphate aldolase class I |
| *pgi* | 0.054 | 5.45E-03 | glucosephosphate isomerase |
| *gpmA* | 0.102 | 2.56E-02 | phosphoglycerate mutase |
| *pfkB* | 0.305 | 1.62E-02 | 6-phosphofructokinase II |
| *gapA* | 0.382 | 2.13E-03 | glyceraldehyde-3-phosphate dehydrogenase A |
| *tpiA* | 0.382 | 8.11E-03 | triosephosphate isomerase |
| *fbaA* | 0.394 | 2.00E-02 | class II fructose-bisphosphate aldolase aldolase |
| *glk* | 0.479 | 1.38E-02 | glucokinase |
| *pykF* | 0.481 | 2.11E-02 | pyruvate kinase I |
| *pgk* | 0.482 | 3.96E-02 | phosphoglycerate kinase |
| *pykA* | 0.500 | 2.33E-03 | pyruvate kinase II |
| **TCA cycle and glyoxylate shunt** | | | |
| *gltA* | 3.681 | 8.52E-05 | citrate synthase |
| *acnA* | 0.270 | 3.33E-02 | aconitate hydratase |
| *icd* | 0.541 | 3.04E-03 | isocitrate dehydrogenase |
| *fumA* | 0.618 | 1.88E-03 | fumarate hydratase |
| *aceA* | 2.348 | 8.73E-04 | isocitrate lyase |
| *aceB* | 2.280 | 8.68E-03 | malate synthase A |
| *aceK* | 1.701 | 1.90E-03 | isocitrate dehydrogenase kinase/phosphatase |
| **Pentose phosphate pathway** | | | |
| *gcd* | 0.062 | 4.49E-03 | glucose dehydrogenase |
| *zwf* | 0.083 | 2.43E-03 | glucose-6-phosphate 1-dehydrogenase |
| *gnd* | 0.168 | 4.91E-03 | 6-phosphogluconate dehydrogenase |
| *alB* | 0.226 | 4.48E-03 | transaldolase B |
| *prs* | 0.305 | 4.79E-02 | phosphoribosylpyrophosphate synthase |
| *gntK* | 0.401 | 3.00E-02 | gluconate kinase 2 |
| **Byproducts (acetate, lactate, formate) metabolism pathway** | | | |
| *acs* | 35844.485 | 1.33E-04 | acetyl-CoA synthetase |

| | | | |
|---|---|---|---|
| *ldhA* | 0.191 | 9.29E-03 | D-lactate dehydrogenase |
| *poxB* | 0.311 | 1.69E-02 | pyruvate dehydrogenase |
| *pflB* | 0.393 | 2.04E-02 | pyruvate formatelyase |

**Supplementary Table 3** Bacteria strains and plasmids used in this study

| Strains and plasmids | Description | Source |
|---|---|---|
| **Strains** | | |
| *E. coli* DH5α | F⁻ *supE*44 Δ*lacU*169 (*ϕ*80 *lacZ* Δ*M15*) *hsdR*17 *recA*1 *endA*1 *gyrA*96 *thi*-1 *relA*1 | Invitrogen |
| *E. coli* χ7213 | *thi*-1 *thr*-1 *leu*B6 *gln*V44 *fhu*A21 *lac*Y1 *rec*A1 *RP4*-2-Tc: Mu *λpir* Δ*asd*A4 Δ*zhf*-2: *Tn*10 | [1] |
| *E. coli* BL21(DE3) | $F^-$ *ompT gal dcm lon hsdSB* ($rB^-$ $mB^-$) *λ*(DE3) | Invitrogen |
| **PG-producing strains** | | |
| Q1944 (WT-PG) | *E. coli* BL21(DE3)/ pA-*accADBC*/ pET-*phlDmar* | [2] |
| Q1963 | *E. coli* BL21(DE3) Δ*arcA* / pA-*accADBC*/ pET-*phlDmar* | This study |
| Q2283 | *E. coli* BL21(DE3) Δ*iclR* / pA-*accADBC*/ pET-*phlDmar* | This study |
| Q1945 | *E. coli* BL21(DE3) $P_{T7}$ *csrB* / pA-*accADBC*/ pET-*phlDmar* | This study |
| Q1964 | *E. coli* BL21(DE3) Δ*ackA* / pA-*accADBC*/ pET-*phlDmar* | This study |
| Q2112 | *E. coli* BL21(DE3)/ pA-*accADBC*/ pET-*phlDmar-acs* | This study |
| Q2176 | *E. coli* BL21(DE3) Δ*arcA*Δ*iclR* / pA-*accADBC*/ pET-*phlDmar* | This study |
| Q1946 | *E. coli* BL21(DE3) Δ*arcA* $P_{T7}$ *csrB* / pA-*accADBC*/ pET-*phlDmar* | This study |
| Q2639 | *E. coli* BL21(DE3) Δ*arcA* Δ*ackA* / pA-*accADBC*/ pET-*phlDmar* | This study |
| Q2113 | *E. coli* BL21(DE3) Δ*arcA* / pA-*accADBC*/ pET-*phlDmar-acs* | This study |
| Q2636 | *E. coli* BL21(DE3) Δ*arcA* Δ*ackA* / pA-*accADBC*/ pET-*phlDmar-acs* | This study |
| Q1947 | *E. coli* BL21(DE3) Δ*arcA* $P_{T7}$ *csrB*Δ*ackA* / pA-*accADBC*/ pET-*phlDmar* | This study |

| | | |
|---|---|---|
| Q2640 | *E. coli* BL21(DE3) Δ*iclR* $P_{T7}$ *csrB* | This study |
| Q2638 | *E. coli* BL21(DE3) Δ*iclR*Δ*ackA* / pA-*accADBC*/ pET-*phlDmar* | This study |
| Q2294 | *E. coli* BL21(DE3) Δ*iclR* / pA-*accADBC*/ pET-*phlDmar-acs* | This study |
| Q2139 | *E. coli* BL21(DE3) $P_{T7}$ *csrB*Δ*ackA* / pA-*accADBC*/ pET-*phlDmar* | This study |
| Q2122 | *E. coli* BL21(DE3) $P_{T7}$ *csrB* / pA-*accADBC*/ pET-*phlDmar-acs* | This study |
| Q2123 | *E. coli* BL21(DE3) Δ*ackA* / pA-*accADBC*/ pET-*phlDmar-acs* | This study |
| Q3753 | *E. coli* BL21(DE3) Δ*arcA* Δ*iclR* $P_{T7}$ *csrB* / pA-*accADBC*/ pET-*phlDmar* | This study |
| Q3754 | *E. coli* BL21(DE3) Δ*arcA* Δ*iclR* Δ*ackA* / pA-*accADBC*/ pET-*phlDmar* | This study |
| Q2641 | *E. coli* BL21(DE3) Δ*iclR* $P_{T7}$ *csrB* Δ*ackA* / pA-*accADBC*/ pET-*phlDmar* | This study |
| Q2637 | *E. coli* BL21(DE3) Δ*iclR* $P_{T7}$ *csrB* / pA-*accADBC*/ pET-*phlDmar-acs* | This study |
| Q2628 | *E. coli* BL21(DE3) Δ*iclR*Δ*ackA* / pA-*accADBC*/ pET-*phlDmar-acs* | This study |
| Q2140 | *E. coli* BL21(DE3) $P_{T7}$ *csrB*Δ*ackA* / pA-*accADBC*/ pET-*phlDmar-acs* | This study |
| Q2627 | *E. coli* BL21(DE3) Δ*iclR* $P_{T7}$ *csrB*Δ*ackA* / pA-*accADBC*/ pET-*phlDmar-acs* | This study |
| Q2956 | *E. coli* BL21(DE3) Δ*arcA* Δ*iclR* $P_{T7}$ *csrB* / pA-*accADBC*/ pET-*phlDmar-acs* | This study |
| Q2957 (ABKS-PG) | *E. coli* BL21(DE3) Δ*arcA* $P_{T7}$ *csrB* Δ*ackA* / pA-*accADBC*/ pET-*phlDmar-acs* | This study |
| Q3752 | *E. coli* BL21(DE3) Δ*arcA* Δ*iclR* $P_{T7}$ *csrB*Δ*ackA* / pA-*accADBC*/ pET-*phlDmar* | This study |
| Q2958 | *E. coli* BL21(DE3) Δ*arcA* Δ*iclR* $P_{T7}$ *csrB*Δ*ackA* / pA- | This study |

| | | |
|---|---|---|
| | *accADBC*/ pET-*phlDmar-acs* | |
| Q2959 | *E. coli* BL21(DE3) Δ*arcA*Δ*iclR*Δ*ackA* / pA-*accADBC*/ pET-*phlDmar-acs* | This study |
| Q4483 | *E. coli* BL21(DE3)/ pET-*phlDmar-aceBA* / pA-*accADBC* | This study |
| Q4484 | *E. coli* BL21(DE3)/ pET-*phlDmar-aceAK6* / pA-*accADBC* | This study |
| **3HP-producing strains** | | |
| Q2191 (WT-3HP) | *E. coli* BL21(DE3)/ pA-*accADBC*/pMCR-N-C | This study |
| Q2709 | *E. coli* BL21(DE3) Δ*arcA* / pA-*accADBC*/pMCR-N-C | This study |
| Q2693 | *E. coli* BL21(DE3) Δ*iclR* / pA-*accADBC*/pMCR-N-C | This study |
| Q2694 | *E. coli* BL21(DE3) $P_{T7}$*csrB* / pA-*accADBC*/pMCR-N-C | This study |
| Q2708 | *E. coli* BL21(DE3) Δ*ackA* / pA-*accADBC*/pMCR-N-C | This study |
| Q2707 | *E. coli* BL21(DE3)/ pA-*accADBC-acs*/pMCR-N-C | This study |
| Q2779 | *E. coli* BL21(DE3) Δ*arcA*Δ*iclR* / pA-*accADBC*/pMCR-N-C | This study |
| Q2780 | *E. coli* BL21(DE3) Δ*arcA* $P_{T7}$*csrB* / pA-*accADBC*/pMCR-N-C | This study |
| Q2781 | *E. coli* BL21(DE3) Δ*arcA* $P_{T7}$*csrB* / pA-*accADBC*/pMCR-N-C | This study |
| Q2782 | *E. coli* BL21(DE3) Δ*arcA* / pA-*accADBC-acs*/pMCR-N-C | This study |
| Q2783 | *E. coli* BL21(DE3) Δ*arcA* $P_{T7}$*csrB* / pA-*accADBC-acs*/pMCR-N-C | This study |
| Q2784 | *E. coli* BL21(DE3) Δ*arcA* $P_{T7}$*csrB*Δ*ackA* / pA-*accADBC*/pMCR-N-C | This study |
| Q2785 | *E. coli* BL21(DE3) Δ*iclR* $P_{T7}$*csrB* / pA-*accADBC*/pMCR-N-C | This study |

| | | |
|---|---|---|
| Q2786 | *E. coli* BL21(DE3) Δ*iclR*Δ*ackA* / pA-*accADBC*/pMCR-N-C | This study |
| Q2787 | *E. coli* BL21(DE3) Δ*iclR* / pA-*accADBC-acs*/pMCR-N-C | This study |
| Q2788 | *E. coli* BL21(DE3) $P_{T7}$*csrB*Δ*ackA* / pA-*accADBC*/pMCR-N-C | This study |
| Q2789 | *E. coli* BL21(DE3) $P_{T7}$*csrB* / pA-*accADBC-acs*/pMCR-N-C | This study |
| Q2790 | *E. coli* BL21(DE3) Δ*ackA* / pA-*accADBC-acs*/pMCR-N-C | This study |
| Q2791 | *E. coli* BL21(DE3) Δ*iclR* $P_{T7}$*csrB*Δ*ackA* / pA-*accADBC*/pMCR-N-C | This study |
| Q3757 | *E. coli* BL21(DE3) Δ*arcA* Δ*iclR* $P_{T7}$*csrB* / pA-*accADBC*/pMCR-N-C | This study |
| Q3758 | *E. coli* BL21(DE3) Δ*arcA* Δ*iclR*Δ*ackA* / pA-*accADBC*/pMCR-N-C | This study |
| Q2697 | *E. coli* BL21(DE3) Δ*iclR* $P_{T7}$*csrB* / pA-*accADBC-acs*/pMCR-N-C | This study |
| Q2696 | *E. coli* BL21(DE3) Δ*iclR*Δ*ackA* / pA-*accADBC-acs*/pMCR-N-C | This study |
| Q2698 | *E. coli* BL21(DE3) $P_{T7}$*csrB*Δ*ackA* / pA-*accADBC-acs*/pMCR-N-C | This study |
| Q2695 | *E. coli* BL21(DE3) Δ*iclR* $P_{T7}$*csrB*Δ*ackA* / pA-*accADBC-acs*/pMCR-N-C | This study |
| Q2908 | *E. coli* BL21(DE3) Δ*arcA*Δ*iclR* $P_{T7}$*csrB* / pA-*accADBC-acs*/pMCR-N-C | This study |
| Q2909 | *E. coli* BL21(DE3) Δ*arcA* $P_{T7}$*csrB*Δ*ackA* / pA-*accADBC-acs*/pMCR-N-C | This study |
| Q2910 | *E. coli* BL21(DE3) Δ*arcA* Δ*iclR* $P_{T7}$ *csrB*Δ*ackA* / pA-*accADBC-acs*/pMCR-N-C | This study |
| Q2911 | *E. coli* BL21(DE3) Δ*arcA*Δ*iclR*Δ*ackA* / pA-*accADBC-acs*/pMCR-N-C | This study |

| | | |
|---|---|---|
| Q3756 | *E. coli* BL21(DE3) Δ*arcA*Δ*iclR* $P_{T7}$*csrB*Δ*ackA* / pA-*accADBC*/pMCR-N-C | This study |
| **glyoxylate-producing strains** | | |
| Q2562 | *E. coli* BL21(DE3)/pETDuet1-*yjhH-xdh-xylC*/ pA-*aldA-yjhG* | [3] |
| Q3769 | *E. coli* BL21(DE3) Δ*arcA* $P_{T7}$*csrB*Δ*ackA*/pETDuet1-*yjhH-xdh-xylC*/ pA-*aldA-acs-yjhG* | This study |
| **LacZ-producing strains** | | |
| Q3833 | *E. coli* BL21(DE3)/pTrcHis2B -*lacZ*/pACYCDuet1 | This study |
| Q3834 | *E. coli* BL21(DE3) Δ*arcA* $P_{T7}$*csrB* Δ*ackA*/ pTrcHis2B -*lacZ*/pACYCDuet1-*acs* | This study |
| **Lysine acetylation research** | | |
| Q3554 | *E. coli* BL21(DE3) / pACYCDuet1/pETDuet1-*gapA* | This study |
| Q3555 | *E. coli* BL21(DE3) / pACYCDuet1/pETDuet1-*pgi* | This study |
| Q3556 | *E. coli* BL21(DE3) / pACYCDuet1/pETDuet1-*pykF* | This study |
| Q3557 | *E. coli* BL21(DE3) / pACYCDuet1/pETDuet1-*glpX* | This study |
| Q3558 | *E. coli* BL21(DE3) / pACYCDuet1/pETDuet1-*zwf* | This study |
| Q3559 | *E. coli* BL21(DE3) / pACYCDuet1/pETDuet1-*gnd* | This study |
| Q3560 | *E. coli* BL21(DE3) / pACYCDuet1/pETDuet1-*aceA* | This study |
| Q3562 | *E. coli* BL21(DE3) / pACYCDuet1/pETDuet1-*icdA* | This study |
| Q3563 | *E. coli* BL21(DE3) / pACYCDuet1/pETDuet1-*mdh* | This study |
| Q3564 | *E. coli* BL21(DE3) / pACYCDuet1/pETDuet1-*poxB* | This study |
| Q3565 | *E. coli* BL21(DE3) / pACYCDuet1/pETDuet1-*ldhA* | This study |
| Q3567 | *E. coli* BL21(DE3) Δ*arcA* $P_{T7}$*csrB* Δ*ackA* /pACYCDuet1-*acs* /pETDuet1-*gapA* | This study |
| Q3568 | *E. coli* BL21(DE3) Δ*arcA* $P_{T7}$*csrB* Δ*ackA* / pACYCDuet1-*acs* /pETDuet-*pgi* | This study |
| Q3569 | *E. coli* BL21(DE3) Δ*arcA* $P_{T7}$*csrB* Δ*ackA* /pACYCDuet1-*acs* /pETDuet1-*pykF* | This study |
| Q3570 | *E. coli* BL21(DE3) Δ*arcA* $P_{T7}$*csrB* Δ*ackA* /pACYCDuet1-*acs* /pETDuet1-*glpX* | This study |
| Q3571 | *E. coli* BL21(DE3) Δ*arcA* $P_{T7}$*csrB* Δ*ackA* / | This study |

| | | |
|---|---|---|
| | pACYCDuet1-*acs* /pETDuet1-*zwf* | |
| Q3572 | *E. coli* BL21(DE3) Δ*arcA* $P_{T7}$*csrB* Δ*ackA* / pACYCDuet1-*acs* /pETDuet1-*gnd* | This study |
| Q3573 | *E. coli* BL21(DE3) Δ*arcA* $P_{T7}$*csrB* Δ*ackA* /pACYCDuet1-*acs* /pETDuet1-*aceA* | This study |
| Q3574 | *E. coli* BL21(DE3) Δ*arcA* $P_{T7}$*csrB* Δ*ackA* / pACYCDuet1-*acs* /pETDuet1-*aceK* | This study |
| Q3575 | *E. coli* BL21(DE3) Δ*arcA* $P_{T7}$*csrB* Δ*ackA* /pACYCDuet1-*acs* /pETDuet1-*icdA* | This study |
| Q3576 | *E. coli* BL21(DE3) Δ*arcA* $P_{T7}$*csrB* Δ*ackA* / pACYCDuet1-*acs* /pETDuet1-*mdh* | This study |
| Q3577 | *E. coli* BL21(DE3) Δ*arcA* $P_{T7}$*csrB* Δ*ackA* / pACYCDuet1-*acs* /pETDuet1-*poxB* | This study |
| Q3578 | *E. coli* BL21(DE3) Δ*arcA* $P_{T7}$*csrB* Δ*ackA* / pACYCDuet1-*acs* /pETDuet1-*ldhA* | This study |
| Q3618 | *E. coli* BL21(DE3) / pACYCDuet1-*acs* | This study |
| Q3619 | *E. coli* BL21(DE3) Δ*arcA* $P_{T7}$*csrB* Δ*ackA* / pACYCDuet1-*acs* | This study |
| **Plasmids** | | |
| pTrcHis2B | $rep_{pBR322}$ $Amp^R$ *lacI*$^q$ $P_{trc}$ | Invitrogen |
| pET30a | $rep_{pBR322}$ $Kan^R$ *lacI*$^q$ $P_{T7}$ | Invitrogen |
| pETDuet1 | $rep_{pBR322}$ $Amp^R$ *lacI* $P_{T7}$ | Invitrogen |
| pACYCDuet1 | $rep_{p15A}$ $Cm^R$ *lacI* $P_{T7}$ | Invitrogen |
| pA-*accADBC* | $rep_{p15A}$ $Cm^R$ *lacI* $P_{T7}$ *accA* $P_{T7}$ *accD* $P_{T7}$ *accBC* | [2] |
| pA-*accADBC-aceA* | $rep_{p15A}$ $Cm^R$ *lacI* $P_{T7}$ *accA* $P_{T7}$ *accD aceA* $P_{T7}$ *accBC* | This study |
| pET-*phlDmar* | $rep_{pBR322}$ $kan^R$ *lacI* $P_{T7}$ *phlD* $P_{T7}$*mar* | [2] |
| pET-*phlDmar-acs* | $rep_{pBR322}$ $kan^R$ *lacI* $P_{T7}$ *phlD* $P_{T7}$*mar acs* | This study |
| pET-*phlDmar-aceAK6* | $rep_{pBR322}$ $kan^R$ *lacI* $P_{T7}$ *phlD* $P_{T7}$*mar aceA aceK*(D477N) | This study |
| pET-*phlDmar-aceBA* | $rep_{pBR322}$ $kan^R$ *lacI* $P_{T7}$ *phlD* $P_{T7}$*mar aceB aceA* | This study |
| pA-*accADBC-acs* | $rep_{p15A}$ $Cm^R$ *lacI* $P_{T7}$ *accA* $P_{T7}$ *accD* $P_{lac,P2-51}$ *acs* $P_{T7}$ *accBC* | This study |

| | | |
|---|---|---|
| pMCR-N-C | rep$_{pBR322}$ Amp$^{R}$ *lacI* P$_{lac,P2-51}$*mcr*$_{1-549}$ P$_{T7}$ *mcr*$_{550-1219}$ (N940V/K1106W/S1114R) | [4] |
| pACYCDuet1-*acs* | rep$_{p15A}$ Cm$^{R}$ *lacI* P$_{T7}$ *acs* | This study |
| pETDuet1-*yjhH-xdh-xylC* | rep$_{pBR322}$ Amp$^{R}$ *lacI* P$_{T7}$ *yjhH xdh* P$_{T7}$ *xylC* | [3] |
| pA-*aldA-yjhG* | rep$_{p15A}$ Cm$^{R}$ *lacI* P$_{T7}$ *aldA* P$_{T7}$ *yjhG* | [3] |
| pA-*aldA-acs-yjhG* | rep$_{p15A}$ Cm$^{R}$ *lacI* P$_{T7}$ *aldA acs* P$_{T7}$ *yjhG* | This study |
| pTrcHis2B -*lacZ* | rep$_{pBR322}$ Amp$^{R}$ *lacI* P$_{Trc}$ *lacZ* | This study |
| pRE112-P$_{T7}$*csrB* | oriT oriV sacB cat P$_{T7}$ *csrB* | This study |
| pETDuet1-*gapA* | rep$_{pBR322}$ Amp$^{R}$ *lacI* P$_{T7}$ *gapA* | This study |
| pETDuet1-*pgi* | rep$_{pBR322}$ Amp$^{R}$ *lacI* P$_{T7}$ *pgi* | This study |
| pETDuet1-*pykF* | rep$_{pBR322}$ Amp$^{R}$ *lacI* P$_{T7}$ *pykF* | This study |
| pETDuet1-*glpX* | rep$_{pBR322}$ Amp$^{R}$ *lacI* P$_{T7}$ *glpX* | This study |
| pETDuet1-*zwf* | rep$_{pBR322}$ Amp$^{R}$ *lacI* P$_{T7}$ *zwf* | This study |
| pETDuet1-*gnd* | rep$_{pBR322}$Amp$^{R}$ *lacI* P$_{T7}$ *gnd* | This study |
| pETDuet1-*aceA* | rep$_{pBR322}$ Amp$^{R}$ *lacI* P$_{T7}$ *aceA* | This study |
| pETDuet1-*icdA* | rep$_{pBR322}$ Amp$^{R}$ *lacI* P$_{T7}$ *icdA* | This study |
| pETDuet1-*mdh* | rep$_{pBR322}$ Amp$^{R}$ *lacI* P$_{T7}$ *mdh* | This study |
| pETDuet1-*poxB* | rep$_{pBR322}$ Amp$^{R}$ *lacI* P$_{T7}$ *poxB* | This study |
| pETDuet1-*ldhA* | rep$_{pBR322}$ Amp$^{R}$ *lacI* P$_{T7}$ *ldhA* | This study |

**Supplementary Table 4** Primers used in this study

| primer | sequence |
|---|---|
| **Chromosomal mutagenesis** | |
| Seq-*Km*-3′ | GGTGAGATGACAGGAGATCC |
| ID-*arcA*-5′ | TCCTGAGGGAAAGTACCCAC |
| ID-*iclR*-5′ | TACGAAATGCCGGATCGTTG |
| ID-*ackA*-5′ | CATAAAACGGATCGCATAACGC |
| $P_{T7}$*csrB*-Up-5′ | CAGAGCTCCAGCTCAGCCAGAATAAGCGCG |
| $P_{T7}$*csrB*-Up-3' | CTATAGGGGAATTGTGAGCGGATAACAATTCCGTCGACA GGGAGTCAGACAAC |
| $P_{T7}$*csrB*-Down-5' | CGCTCACAATTCCCCTATAGTGAGTCGTATTAGAAGATAG AATCGTCTTTTTC |
| $P_{T7}$*csrB*-Down-3' | ATCTGCGGTACCGTGGCATGAAGAGCATAAAA |
| ID-*csrB*-5' | TTCCAGCATTAGCTCGCATC |
| $P_{T7}$ promoter-3′ | TTGTTATCCGCTCACAATTC |
| **Plasmids for fermentation** | |
| pTrcHis2B-*lacZ*-5′ | CATGCCATGGCCATGATTACGGATTCAC |
| pTrcHis2B-*lacZ*-3′ | CAGTGTCGACTTATTTTTGACACCAGACCAAC |
| pACYCDuet1-*acs*-5′ | CATGCCATGGGCATGAGCCAAATTCACAAACAC |
| pACYCDuet1-*acs*-3′ | CCGGAGCTC TTACGATGGCATCGCGATAGC |
| pA-*accADBC-acs*-5′(overlap-1) | GTTGTGTGGAAGGGGAATTGTGAGCGGATAACAATTCCC CTGTCACATATTATTAACATCCTA |
| pA-*accADBC-acs*-5′(overlap-2) | CCGCTTAAGCTTTACACTTTAAGCTTCATATGTTTATGTTG TGTGGAAGGGGAATTG |
| pA-*accADBC-acs*-3′ | ACGCGTCGACAAAAGCCTCCGGTCGGAGGCTTTTTTACG ATGGCATCGCGATAG |
| pA-*aldA-acs-yjhG*-5′ | CCGGAATTCCCTACATTTAACGCTTATGC |
| pA-*aldA-acs-yjhG*-3′ | CCCAAGCTTTTACGATGGCATCGCGATAGC |
| pET-*phlDmar-acs*-5′ | ACGCGTCGACCACATATTATTAACATCCTA |
| pET-*phlDmar-acs*-3′ | CCGAAGCTTTTACGATGGCATCGCGATAG |
| pET-*phlDmar-aceBA*-5′ | CCGAAGCTTGAAACGTACCTCAGCAGGTG |
| pET-*phlDmar-aceBA*-3′ | CCGCTCGAGTTAGAACTGCGATTCTTCGG |

| | |
|---|---|
| pET-*phlDmar-aceAK6*-Up-5′ | CCGAAGCTTCGATGCCGCACGCTTGATGG |
| pET-*phlDmar-aceBA*-Up -3′ | CCGTCATGTAGCAAATTTCATTGTAATCATAAAAAACCACAC |
| pET-*phlDmar-aceAK6*-Down-5′ | GTGTGGTTTTTTATGATTACAATGAAATTTGCTACATGACGG |
| pET-*phlDmar-aceBA*-Down-3′ | CCGCTCGAGTCAAAAAGCATCTCCCCATAC |
| **Plasmids for acetylation and phosphorylation analysis** | |
| pETDuet1-*gapA*-5′ | CCGGGATCCGATGACTATCAAAGTAGGTATC |
| pETDuet1-*gapA*-3′ | CCGGAGCTCTTATTTGGAGATGTGAGCGA |
| pETDuet1-*pgi*-5′ | CCGGGATCCG ATGAAAAACATCAATCCAACG |
| pETDuet1-*pgi*-3′ | CCGGAGCTCTTAACCGCGCCACGCTTTATAG |
| pETDuet1-*pykF*-5′ | CCGGGATCCGATGAAAAAGACCAAAATTGTTTG |
| pETDuet1-*pykF*-3′ | CCGGAGCTCTTACAGGACGTGAACAGATG |
| pETDuet1-*glpX*-5′ | CCGGGATCCGATGAGACGAGAACTTGCCATC |
| pETDuet1-*glpX*-3′ | CCGGAGCTCTCAGAGGATGTGCACCTGCA |
| pETDuet1-*zwf*-5′ | CCGGGATCCGATGGCGGTAACGCAAACAGC |
| pETDuet1-*zwf*-3′ | CCGGAGCTCTTACTCAAACTCATTCCAGG |
| pETDuet1-*gnd*-5′ | CCGGGATCCGATGTCCAAGCAACAGATCGG |
| pETDuet1-*gnd*-3′ | CCGGAGCTCTTAATCCAGCCATTCGGTATG |
| pETDuet1-*aceA*-5′ | CCGGGATCCGATGAAAACCCGTACACAACA |
| pETDuet1-*aceA*-3′ | CCGGAGCTCTTAGAACTGCGATTCTTCAG |
| pETDuet1-*icdA*-5′ | CCGGGATCCGATGGAAAGTAAAGTAGTTGT |
| pETDuet1-*icdA*-3′ | CCGGAGCTCTTACATGTTTTCGATGATCG |
| pETDuet1-*mdh*-5′ | CCGGGATCCGATGAAAGTCGCAGTCCTCGG |
| pETDuet1-*mdh*-3′ | CCGGAGCTCTTACTTATTAACGAACTCTTC |
| pETDuet1-*poxB*-5′ | CCGGGATCCGATGAAACAAACGGTTGCAGC |
| pETDuet1-*poxB*-3′ | CCGGAGCTCTTACCTTAGCCAGTTTGTTTTC |
| pETDuet1-*ldhA*-5′ | CCGGGATCCGATGAAACTCGCCGTTTATAGC |
| pETDuet1-*ldhA*-3′ | CCGGAGCTCTTAAACCAGTTCGTTCGGGC |
| **qRT-PCR** | |

| | |
|---|---|
| qPCR-*patZ*-5′ | CAGGTTACCTGATGATGCGT |
| qPCR-*patZ*-3′ | CGCAAGGTCGGGTGTAAAGG |
| qPCR-*cobB*-5′ | GCGAGCGTTTGCGTCAGCGT |
| qPCR-*cobB*- 3′ | CAACCCGATGTTCTTCCCAC |

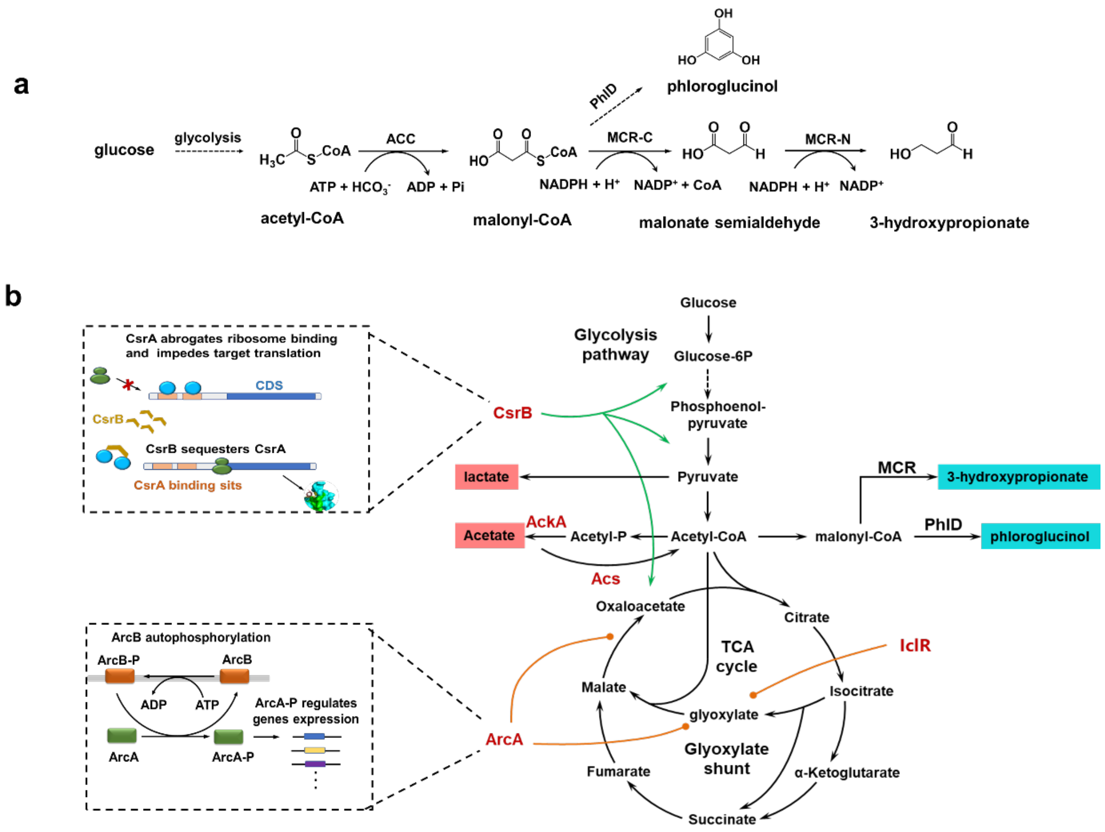


**Supplementary Fig. 1** Genes modified in this study and their functions. (a) Phloroglucinol and 3-hydroxypropionate biosynthetic pathways from glucose. ACC, acetyl-CoA carboxylase; MCR-N and MCR-C, N- and C-terminal fragments of malonyl-CoA reductase; PhlD, polyketide synthase. (b) The regulatory functions or catalyzed reactions of genes modified in this study.

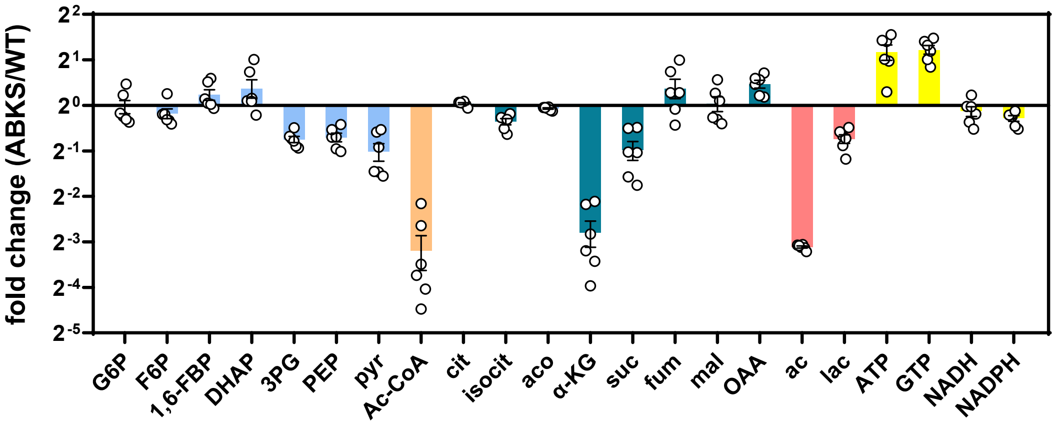


**Supplementary Fig. 2** Changes of intracellular metabolites in WT-PG and ABKS-PG strains ($n$ = 6 biologically independent samples). Colors represent the categories of corresponding metabolites. Blue, glycolysis intermediate; green, intermediate of TCA cycle; red, byproduct; yellow, energy and reducing power.

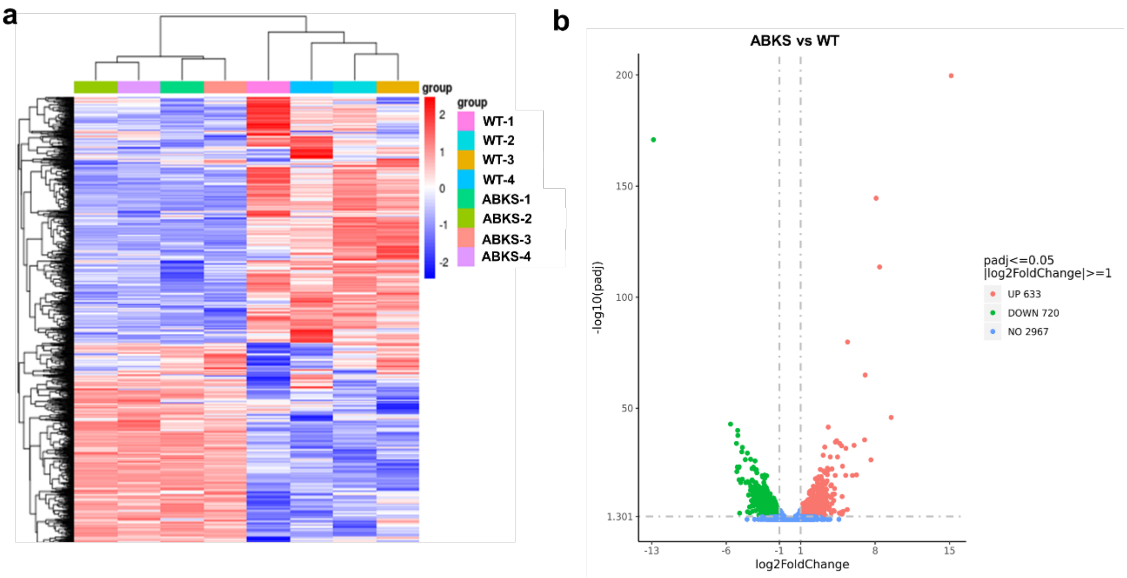


**Supplementary Fig. 3** Overview of transcriptome analysis of WT-PG and ABKS-PG strains. (a) Hierarchical clustering analysis of the differentially expressed genes (DEGs) between those two strains, (b) The overall distribution of DEGs. The smaller corrected p-Value (padj) represents more significant difference.

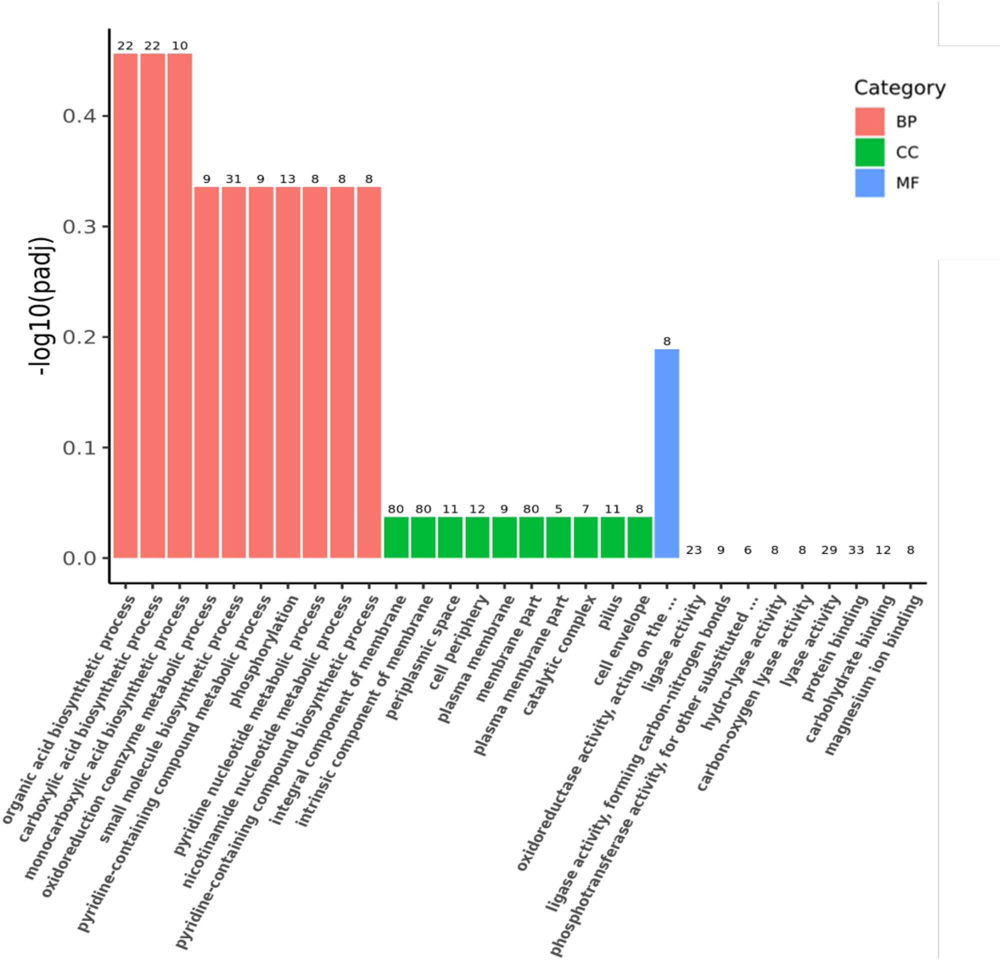


**Supplementary Fig. 4** Gene Ontology (GO) analysis of DEGs. BP, biological processes; MF, molecular functions; CC, cellular components. The number of different genes was listed on the bar. The smaller corrected p-Value (padj) represented more significant difference.

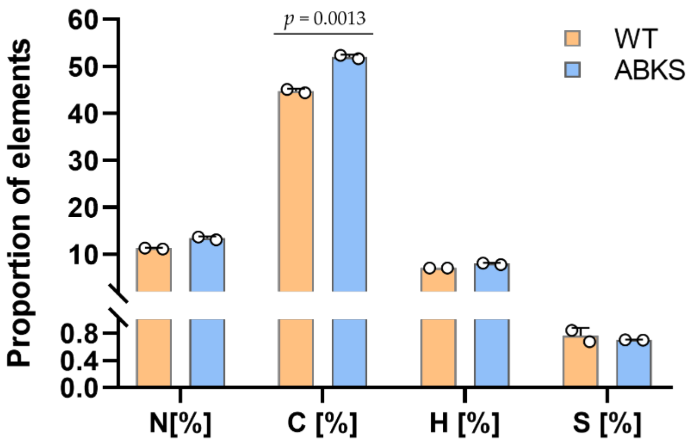


**Supplementary Fig. 5** The proportion of N, C, H, S element of WT-PG and ABKS-PG strains ($n$ = 2 biologically independent samples). Error bars, mean ± SEM.

Supplementary references:

1. Roland K, Curtiss R, 3rd, Sizemore D. Construction and evaluation of a D*cya* D*crp Salmonella typhimurium* strain expressing avian pathogenic *Escherichia coli* O78 LPS as a vaccine to prevent airsacculitis in chickens. *Avian Dis* **43**, 429-441 (1999).

2. Cao YJ, Jiang XL, Zhang RB, Xian M. Improved phloroglucinol production by metabolically engineered *Escherichia coli*. *Appl Microbiol Biotechnol* **91**, 1545-1552 (2011).

3. Liu M, Ding Y, Xian M, Zhao G. Metabolic engineering of a xylose pathway for biotechnological production of glycolate in Escherichia coli. *Microb Cell Fact* **17**, 51 (2018).

4. Liu C, Ding Y, Zhang R, Liu H, Xian M, Zhao G. Functional balance between enzymes in malonyl-CoA pathway for 3-hydroxypropionate biosynthesis. *Metab Eng* **34**, 104-111 (2016).
